# Pseudolysogeny-mediated evolutionary trade-offs favor phage therapy by limiting antibiotic resistance and virulence in *Cutibacterium acnes*

**DOI:** 10.1101/2025.09.15.676368

**Authors:** Abigail Trejo-Hernández, Alberto Checa, Patricia Sepúlveda-Gutíerrez, Andrés Andrade-Domínguez

**Affiliations:** Eubiology Labs, Research and Development, Morelos, México; Dermatoss Vanguardia Médica Clinic, Morelos, México

## Abstract

Phage therapy has mostly focused on strictly lytic phages, yet the ecological and evolutionary implications of pseudolysogeny remain poorly understood. Pseudolysogeny—where a phage genome persists in a non-integrated, latent state within the host—has been largely overlooked due to concerns about therapeutic efficacy. Here, we demonstrate that pseudolysogeny in *Cutibacterium acnes* confers superinfection resistance but imposes substantial fitness costs, including reduced biofilm formation, impaired interspecies competitiveness, and reversal of antibiotic resistance. Pseudolysogenic phages were also capable of killing starved *C. acnes* cells through direct lytic replication. No evidence of transduction of clindamycin resistance by these phages was detected. In a 3-month proof-of-concept study, topical application of a pseudolysogenic phage significantly reduced *C. acnes* abundance, acne lesions, and inflammation, with no adverse effects and persistence of viable phages post-treatment. Importantly, no phage-resistant *C. acnes* clones were detected during the *in vivo* study, likely due to the evolutionary trade-offs associated with pseudolysogeny that diminish the bacterial ecological fitness. These findings highlight pseudolysogeny as a mechanism that can be leveraged to enhance phage therapy outcomes while maintaining microbiome stability and limiting antibiotic resistance evolution.

## Introduction

Acne affects up to 85% of adolescents and a growing number of adults, with significant health impacts 1,2. Its pathogenesis is linked to the overgrowth of *C. acnes* 3. Current treatments, such as antibiotics, retinoids, and benzoyl peroxide, face limitations like adverse effects, poor long-term efficacy, and antibiotic resistance 4–6. These limitations underscore the need for alternative strategies that preserve commensal skin microbiota 7 6.

Phage therapy has resurfaced as a promising antibacterial approach, offering host specificity and reduced disruption of commensal microbiota compared to conventional antibiotics8,9 . Phages targeting *C. acnes* are abundant in the human skin microbiome, and studies have shown a reduced prevalence of phages in acne-affected individuals, suggesting a natural role in controlling *C. acnes* populations10,11. Moreover, *C. acnes*-specific phages have demonstrated *in vitro* and *in vivo* efficacy in lysing pathogenic strains and reducing acne-associated inflammation12,13. Despite this potential, a comprehensive understanding of the ecological and evolutionary dynamics of phage–host interactions—including the emergence of phage resistance and microbiome interplay—is essential prior to clinical application.

A critical but underexplored aspect of *C. acnes* phage biology is pseudolysogeny. Unlike classical lysogeny, *C. acnes* phages lack integrases and do not integrate into the bacterial genome14–16. Instead, many *C. acnes* phages can persist as episomal DNA in a pseudolysogenic state, where they do not immediately lyse the host nor stably integrate, but can influence bacterial physiology14. This pseudolysogenic state induces superinfection exclusion (SIE), whereby infected bacteria become resistant to subsequent phage attacks via expression of phage-encoded proteins17. Although pseudolysogeny may limit the immediate bactericidal effect of phage therapy, its broader ecological and evolutionary consequences remain poorly understood.

Here, we investigated the ecological and evolutionary dynamics, efficacy, and safety of pseudolysogenic phage therapy against *C. acnes* through a human-based clinical proof-of-concept trial. We evaluated: (i) the clinical impact of phage therapy on the reduction of acne lesions, erythema, and inflammation; (ii) its effects on *C. acnes* and *Staphylococcus* spp., key members of the skin microbiome; and (iii) the emergence of phage-resistant strains and the potential fitness costs associated with pseudolysogeny. In addition, we examined whether the CaCom2 phage could mediate horizontal gene transfer of macrolide resistance, a clinically significant concern in the treatment of *C. acnes* infections.

Our results demonstrate that pseudolysogenic phage therapy not only reduces *C. acnes* abundance and improves clinical outcomes in acne vulgaris, but also imposes strong evolutionary trade-offs on the pathogen. These trade-offs limit biofilm formation, reduce competitive fitness, and even reverse antibiotic resistance. Notably, no phage-resistant *C. acnes* clones emerged during treatment, and no horizontal gene transfer of resistance determinants was detected. Together, these findings position pseudolysogeny as an eco-evolutionary constraint that can be harnessed in phage therapy to suppress resistance emergence and improve long-term therapeutic efficacy.

## Material and methods

### Isolation of *Cutibacterium* species from acne lesions

Facial sebum samples were collected from 21 individuals to isolate *Cutibacterium spp.* First, the sampling area was cleaned with a sterile cotton swab moistened in 70% ethanol. Then, a sterile bacteriological loop was used to swab the infected sebaceous follicles, and the samples were immediately streaked onto BD BACTO™ Brain Heart Infusion agar plates supplemented with 5% sheep blood. These plates were placed in a sealed container with a BD GasPak™ EZ Anaerobe Container System sachet and incubated at 36°C for seven days. Identification of *C. acnes* by PCR <u>18</u> and Nanopore sequencing of the 16S rRNA gene for bacterial identification are described in Supplementary Material.

### Antibiotic Susceptibility and genetic resistance testing

Isolates were tested for susceptibility to ampicillin, tetracycline, minocycline, doxycycline, erythromycin, and clindamycin using disc diffusion on MHA under anaerobic conditions, following CLSI guidelines. Clindamycin resistance genes *erm(X)* and *erm(50)* were detected by PCR. Full protocols are provided in the Supplementary Material.

### Isolation and selection of phages for acne phage therapy

Eight phages were isolated from sebum samples of acne-free adolescents and propagated using *C. acnes* 11827. Detailed protocols for phage isolation, amplification, host range determination, and thermal stability are provided in the Supplementary Materials and Methods online.

### Isolation of *C. acnes* pseudolysogens resistant to phage **Φ**CaCom2

ΦCaCom2-resistant pseudolysogens were isolated by repeated exposure of *C. acnes* 11827 lawns to high-titer phage lysate (7×10^10^ PFU/mL), followed by selective re-isolation from lysis zones. Phage-resistant colonies, in which the presence of phage DNA was confirmed, were subjected to investigation in order to ascertain the presence of the circular phage genome within the cells using a PCR approach employing primers that specifically bind to the ends of the phage genomes (CaEnd-F: 5′-CCGAAGCCGACCACATCACACC-3′, CaEnd-R: 5′-TCATCCAACACCTGCTGCTGCC-3′)14. Full protocols are detailed in the Supplementary Material.

### Transduction experiments

A total of 10 mL of a C. acnes 11827 recipient culture (5 × 10 CFU/mL) was infected with ΦCaCom2 phage propagated on the clindamycin-resistant donor strain YIJA, to reach a final phage concentration of 5 × 10 PFU/mL. After 24 hours of incubation, total viable cells were quantified by plating serial dilutions on non-selective BHI agar. To assess transduction of clindamycin resistance, 1 mL of the culture was plated on BHI agar supplemented with 30 µg/mL clindamycin. The number of pseudolysogens was estimated by enumerating phage-resistant colonies, which were subsequently verified by colony PCR.

### Assessment of pseudolysogen fitness in monoculture and co-culture conditions

The fitness of pseudolysogens Ps-1 and Ps-2 (derived from *C. acnes* 11827) was evaluated in monoculture and co-culture with *C. avidum*, *C. granulosum*, or *S. epidermidis* in heat-inactivated fetal bovine serum under anaerobic conditions. Fitness was calculated using the Malthusian parameter (m = ln(Nf/Ni)/t), where *Nf* and *Ni* are the final and initial CFU/mL, and *t* is time in days, following Lenski *et al*.19. Full details are available in the Supplementary Materials and Methods online.

### Phage production and quality control for therapeutic use

Phage production was carried out by the Phage Therapy Unit at CAS Biotechnology, a facility licensed by COFEPRIS (permit no. 23 1701 5018 X0080), following procedures described in the Supplemental Material and Methods online. Production followed the “Monograph for Phages APIS version 1” developed by the FAMHP and the Belgian Scientific Institute of Public Health 20, ensuring compliance with quality standards for magistral phage preparations. Topical formulations were prepared in accordance with the Mexican Good Manufacturing Practices for Cosmetics (NOM-259-SSA1-2022) 21.

The final phage stock was tested for sterility (aerobic and anaerobic bacteria, fungi) and cytotoxicity in HaCat cells. The formulation was approved by the CAS Biotechnology health officer for use in acne volunteers.

### Phage **Φ**CaCom2 topical solution: composition and treatment regimen

The phage ΦCaCom2 was formulated in a topical solution, which was administered by spraying (∼200 µL) onto the face every 24 hours during the night. The formulation name PB7L is composed of distilled water, 100 mM NaCl, 40 mM Tris pH 7.5, 10 mM MgSO4, a mixture of stabilizing agents for phages, and ∼5e8 UFP/mL of phage ΦCaCom2. The solution was packaged in sterile amber-colored glass vials. Each participant was provided with 4 vials, each containing 30 mL, which was sufficient for a 3-month treatment. The participants stored the vials at room temperature between 25-34°C, avoiding exposure to temperatures higher than 37°C and direct sunlight exposure.

### Ethical considerations and informed consent

This proof-of-concept study was conducted in accordance with the Declaration of Helsinki (2024 revision)22 and relevant Mexican regulations, including the Reglamento de la Ley General de Salud en Materia de Investigación para la Salud23, which governs minimal-risk, non-interventional studies. As such, formal ethics committee approval was not required.

Participants received a topical formulation containing bacteriophages and cosmetic-grade ingredients. The bacteriophages comply with Mexican and international safety regulations, including their recognition as food additives and processing aids under the Acuerdo de aditivos y coadyuvantes (DOF 16/07/2012)24, and are compatible with NOM-141-SSA1/SCFI-2012 for cosmetic products25.

Subjects had a documented history of unsuccessful responses to conventional acne treatments and provided written informed consent after being fully informed of the study’s objectives and the nature of the investigational product. Non-invasive procedures or collection of sensitive personal data were involved. All clinical information was anonymized to ensure participant confidentiality. The study was internally reviewed and classified as ethically exempt. Informed consent was obtained for the publication of anonymized information and any identifiable images in an open-access scientific journal.

### Study design and clinical outcome assessments

Patients were monitored at baseline, after the first week, and subsequently at two-week intervals throughout the 12-week treatment period and during the follow-up phase. Clinical evaluations were performed by two independent dermatologists (Dr. Patricia Sepúlveda Gutiérrez and Dr. Gissel Castellanos Ramos) and included quantification of inflammatory and non-inflammatory acne lesions, as well as severity assessments using the Cunliffe classification system. Adverse events were recorded at each follow-up visit.

### Eligibility and study endpoints

Adults (≥18 years) with mild to moderate facial acne and confirmed *C. acnes* colonization were eligible. Key exclusion criteria included recent use of acne treatments or other bacterial colonization. Co-primary endpoints at Week 8 included inflammatory lesion count reduction and IGA score improvement. Safety assessments included treatment-emergent adverse events (TEAE), vital signs, physical examinations, and site tolerability. Patient-reported outcomes included acne severity and discomfort scores on visual and numerical analog scales. Full inclusion criteria and assessment details are provided in the Supplementary Material.

### Sampling of the cultivable microbiome

Facial skin microbiota was sampled from standardized 1 cm² areas using sterile swabs, targeting both comedonal and papulopustular lesions. Samples were processed immediately, cultured on BHI blood agar (anaerobically, 8 days) and Mannitol salt agar (MSA) plates (aerobically, 72 h) for quantification and isolation of key bacterial taxa. Full sampling and processing procedures are detailed in the Supplementary Material.

### Statistical analysis

Statistical analyses and data visualization were performed using GraphPad Prism version 8.0.2 (GraphPad Software, La Jolla, CA). Data are presented as meanL±Lstandard deviation (SD). Normality, homogeneity of variance, and specific statistical tests are detailed in the figure legends. Exact *p*-values, effect sizes, and confidence intervals are also provided in the legends. All experiments were conducted with at least three independent biological replicates, unless stated otherwise.

## Results

### Selection of phages for therapy against *acne vulgaris*

Eight phages were isolated from the facial skin of healthy children and propagated using *C. acnes* ATCC 11827. Their lytic activity was tested against 22 clinical *C. acnes* isolates and two laboratory strains to identify candidates with broad therapeutic potential. Phages ΦCaCom2, ΦINE1, and ΦCaSA-2 lysed all *C. acnes* strains tested but showed no activity against *C. granulosum* or *C. avidum*. This indicates that these phages have a broad host range among *C. acnes* clinical isolates but are unable to infect other species within the *Cutibacterium* genus (Table 1) 26,27.

**Table 1.**
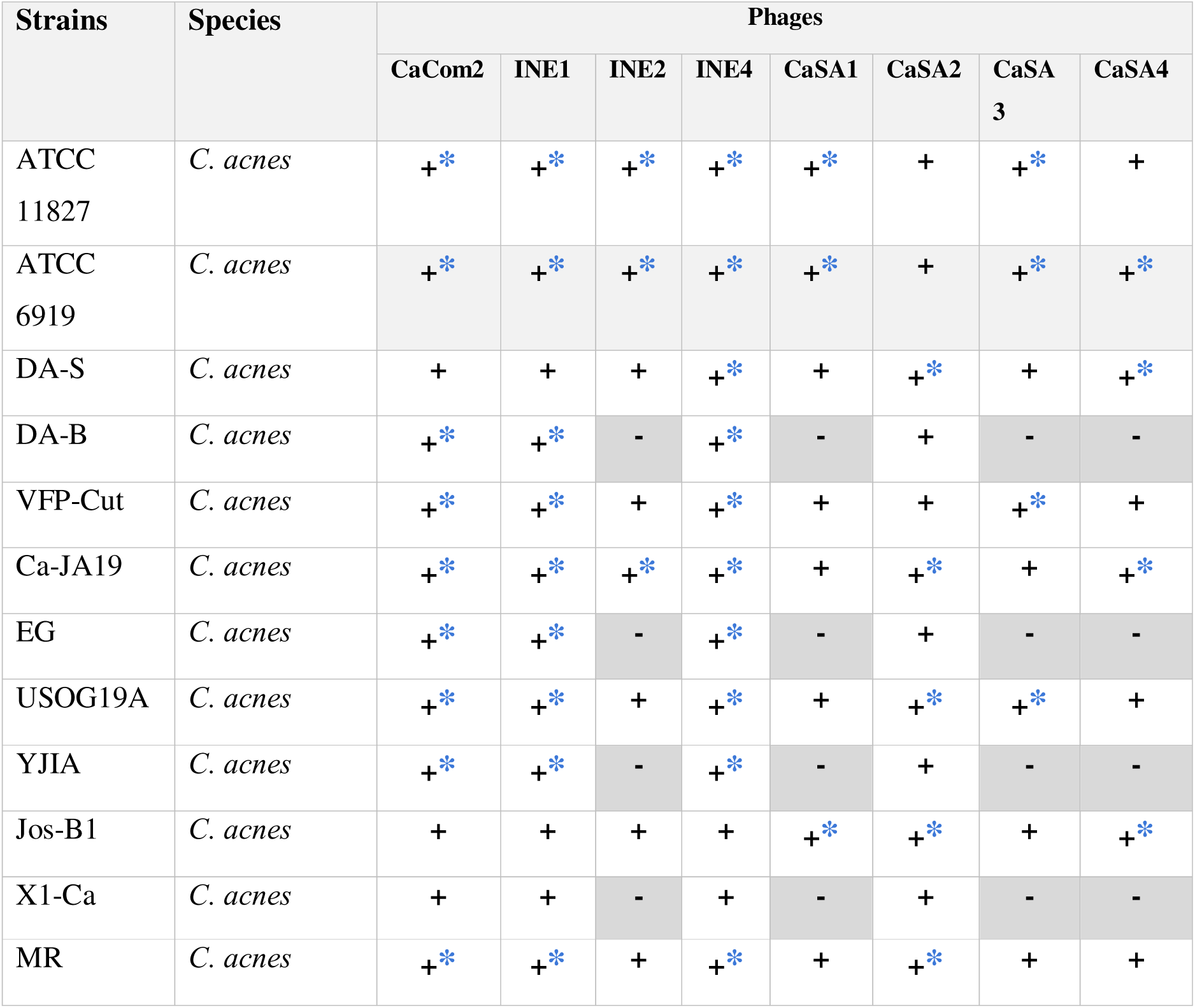
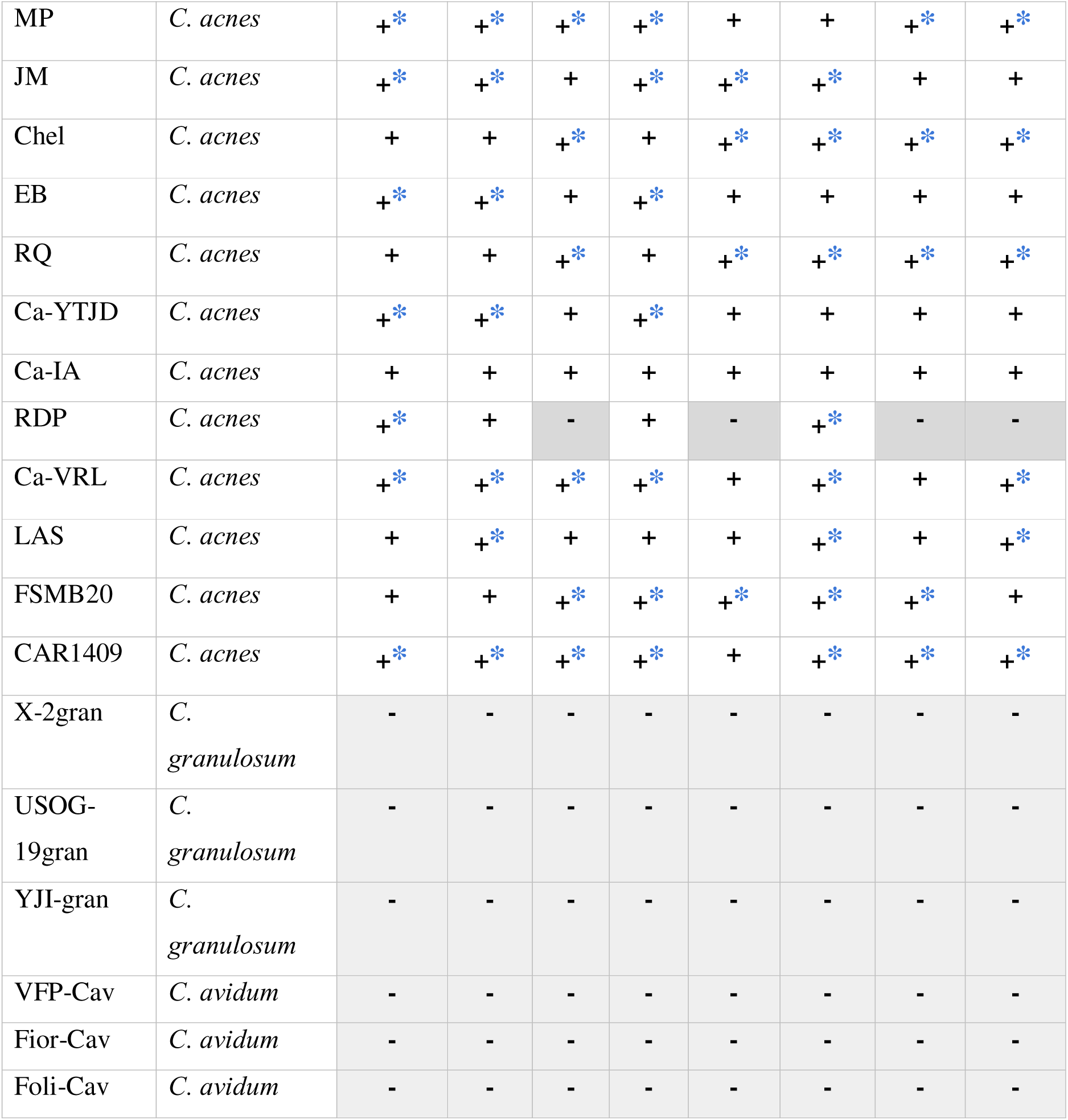
Host Range of *C. acnes* Phages. This table summarizes the host range of phages tested against *C. acnes*, *C. granulosum*, and *C. avidum* strains. Susceptibility was assessed based on the ability of each phage to form lysis plaques in double-layer agar assays. Lysis is indicated by “+”, while resistance is denoted by “−”. * indicates the establishment of pseudolysogeny.

ΦCaCom2 was ultimately selected for therapeutic use based on additional criteria beyond host range. This phage exhibited superior thermal stability compared to ΦINE1 and ΦCaSA-2, with only a 1.5-log reduction in titer after 12 months at room temperature (24–33 °C) and under simulated transport conditions (37–42 °C for 12 hours) (see Supplementary Fig. S1 online). Its high stability, broad host range, and ability to disrupt biofilms supported the selection of ΦCaCom2 for application in the clinical phase of the study.

### The **Φ**CaCom2 phage suppresses growth in liquid culture and disrupts biofilm of clindamycin-resistant isolates

Analysis of *C. acnes* growth kinetics in agitated liquid culture revealed that the ΦCaCom2 phage, either alone or as part of a cocktail (ΦCaCom2, ΦINE1, ΦCaSA2), significantly outperformed 30 µg/mL clindamycin in suppressing the growth of strain 11827, which is sensitive to this antibiotic (Fig.L1A).

Among the 22 clinical isolates obtained from acne patients, 9 were resistant to 30 µg/mL clindamycin (Supplementary TableLS1) and exhibited an enhanced capacity for biofilm formation compared to the reference strain 11827 (see Supplementary Fig.LS2 online). Biofilms were established during the early stages of growth on polystyrene plates and served as a model to assess phage-mediated biofilm disruption. Despite all isolates being susceptible to ampicillin (Supplementary TableLS1), this antibiotic had no observable effect on biofilm biomass (Fig.L1B). In contrast, the ΦCaCom2 phage alone was as effective as the phage cocktail in eliminating and reducing biofilms formed by all clindamycin-resistant isolates (Fig.L1B). Furthermore, both phage treatments were significantly more effective than clindamycin or ampicillin in dispersing established biofilms.

### ΦCaCom2 replicates in stationary-phase *C. acnes* population, overcoming a common limitation of phage therapy

It has previously been widely reported that bacteria in the stationary phase or under nutrient-limited conditions exhibit increased tolerance to both antibiotics and phage infection. This resistance is often attributed to reduced metabolic activity and limited availability of host replication machinery, which impair phage adsorption and propagation. In particular, Maffei *et al*.28 demonstrated that most phages fail to replicate in stationary-phase *Pseudomonas aeruginosa* cells, both in vitro and in vivo, highlighting a major limitation for phage therapy in non-dividing bacterial populations.

Interestingly, our results indicate that ΦCaCom2 can actively replicate within stationary phase populations of *C. acnes* for up to seven days. While treatment with 10 µg/mL ampicillin had no significant impact on cell viability over a 12-day period, infection with ΦCaCom2 resulted in a two-log reduction in CFU/mL and a concurrent three-log increase in PFU/mL, demonstrating active phage replication (Fig. 2A–B). In contrast, the ΦCaSa2 phage, which shares the same host range as the ΦCaCom2 phage, did not show a statistically significant reduction in viability or an increase in the concentration of free phages (Fig. 2A-B). Furthermore, ΦCaCom2 produced visible lysis zones in high-density starved cells pre-grown for 7 days on BHI agar, providing additional evidence of its lytic activity under conditions that typically impair phage efficacy (Fig. 2C).

**Fig. 1.**
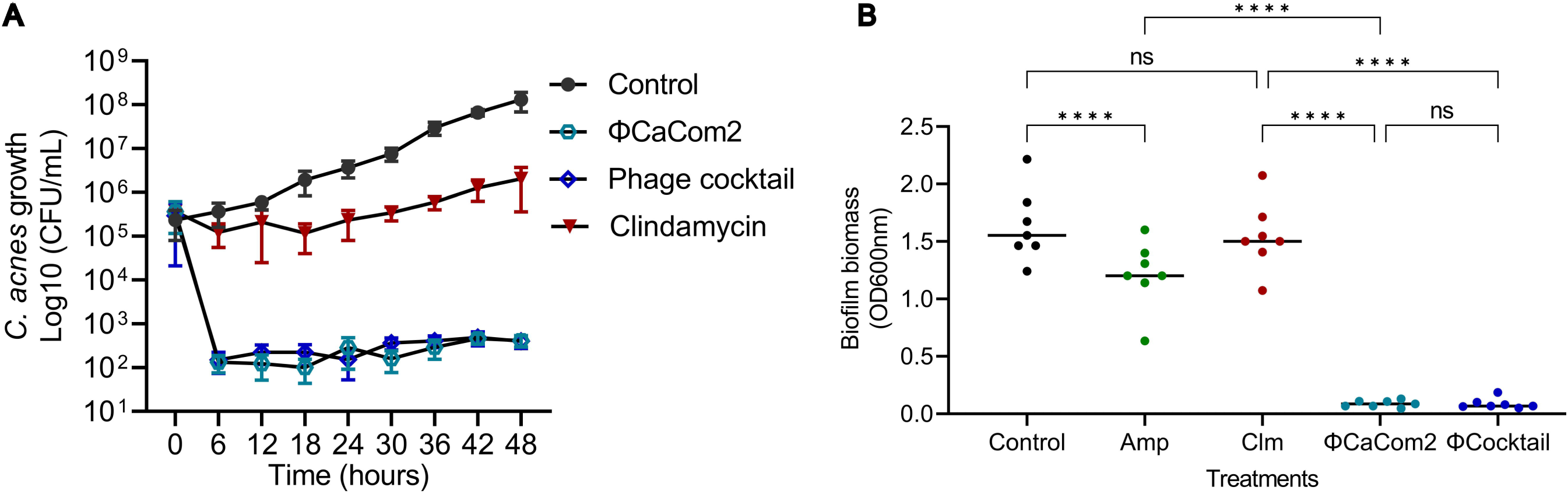
Φ**CaCom2 suppresses growth and biofilm formation of clindamycin-resistant *C. acnes*. (A)** Bacterial growth curves show that ΦCaCom2 (3L×L10L PFU/mL) alone is as effective as a phage cocktail (ΦCaCom2, ΦINE1, ΦCaSA2; each at 1L×L10L PFU/mL) in suppressing *C. acnes* 11827 growth in liquid culture. Both treatments outperformed clindamycin (30Lµg/mL). Statistical analysis of AUC values using Kruskal–Wallis followed by Dunn’s multiple comparisons test showed that ΦCaCom2 and the phage cocktail significantly outperformed clindamycin (***pL<L0.0001 and *pL=L0.0004, respectively), while no significant difference was observed between ΦCaCom2 and the phage cocktail. Data are meanL±Ls.d. from three independent experiments (*n*L=L3). **(B)** Biofilm biomass quantified after 72Lh of incubation using crystal violet staining. Treatments with ΦCaCom2 (3L×L10LLPFU/mL), ΦINE1 (3L×L10LLPFU/mL), and a phage cocktail (ΦCaCom2, ΦINE1, ΦCaSA2; each at 1L×L10LLPFU/mL) were tested against seven clindamycin-resistant *C. acnes* isolates and compared to ampicillin (50Lµg/mL) and clindamycin (30Lµg/mL). Each data point represents the mean biomass of each isolate from three independent experiments. Two-way ANOVA revealed significant main effects of treatment (*p*L<L0.0001) and isolate (*p*L=L0.0001), as well as a significant treatment–isolate interaction (*p*L<L0.0001). Tukey’s multiple comparisons test showed that ΦCaCom2 and the phage cocktail significantly reduced biofilms biomass compared to both antibiotics (**** *p*L<L0.0001). No significant difference (*ns*) was observed between ΦCaCom2 and the phage cocktail (*p*L=L0.99).

**Figure 2.**
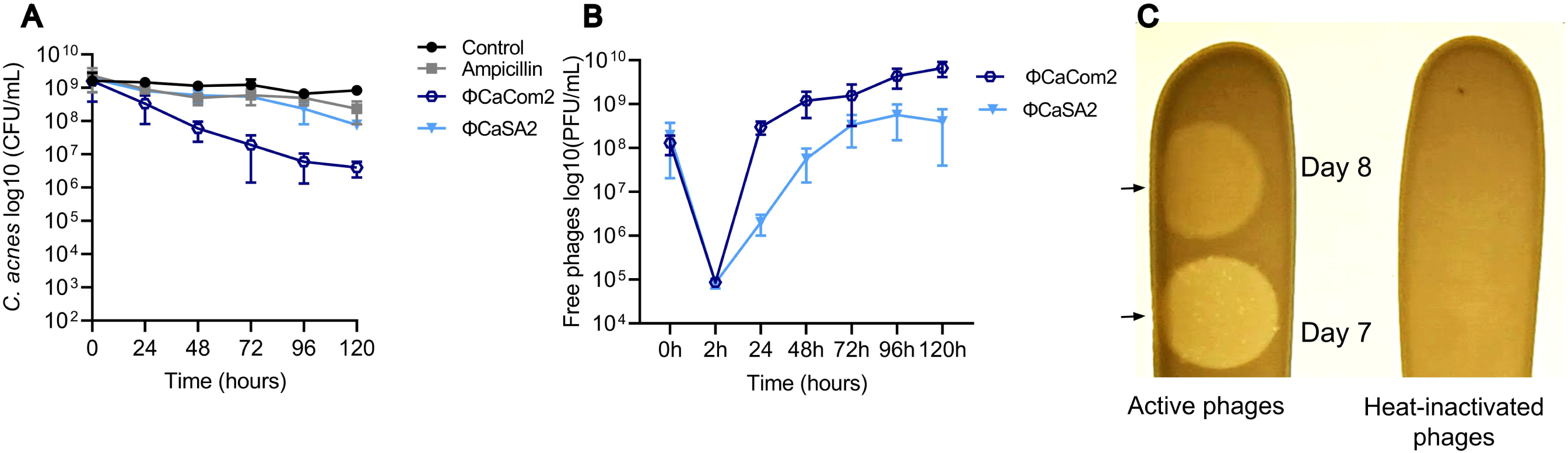
**Φ**CaCom2 outperforms antibiotics in reducing viability and replicates efficiently in stationary-phase *C. acnes*. (**A**) To evaluate lytic activity under late stationary-phase in brain heart infusion (BHI) broth, the viability of *C. acnes* cells (CFU/mL) from a 7-day-old culture was monitored following exposure to ΦCaCom2 (1L×L10L PFU/mL), ΦCaSa2 (1L×L10L PFU/mL), 10Lµg/mL ampicillin, or heat-inactivated phage lysate (control). Viable cell curves analyzed by repeated-measures two-way ANOVA with Tukey’s post hoc test revealed significant differences: ΦCaCom2 vs control (*p*L=L0.021), ΦCaCom2 vs ampicillin (*p*L=L0.0357), ΦCaSA2 vs control (*p*L=L0.048). No significant differences were found for control vs ampicillin (*ns*), ΦCaCom2 vs ΦCaSA2 (*p*L=L0.149), or ampicillin vs ΦCaSA2 (*p*L=L0.7442). Data are meanL±Ls.d. from three independent experiments (*n*L=L3). (**B**) Phage replication was assessed by quantifying phage titers at 30 minutes and then every 24 hours after infection of the 7-day-old culture with ΦCaCom2 (1L×L10L PFU/mL) and ΦCaSa2 (1L×L10L PFU/mL). Statistically significant differences between the ΦCaCom2 and ΦCaSa2 replication curves were detected at 24 and 48 hours (repeated-measures two-way ANOVA followed by Bonferroni’s multiple comparisons test, *p* < 0.05). Data are meanL±Ls.d. from three independent experiments (*n*L=L3). (**C**) Representative image from three independent experiments showing a lysis zone (arrows) on *C. acnes* lawns caused by 5LµL of ΦCaCom2 (2L×L10L PFU/mL) on days 7 and 8. Heat-inactivated lysates served as negative controls. All assays were performed on cultures grown in BHI, and images were captured 24Lh post-infection.

One possible explanation for this uncommon replicative behavior lies in the metabolic heterogeneity of bacterial populations. It has been proposed that bacterial populations in aging or starved colonies are composed of subpopulations of dormant, apoptotic, and metabolically active cells, with the latter relying on nutrients released by the death of their neighbors 29 30,31,32. We hypothesize that phage infection in these nutrient-limited environments initiates lysis in the metabolically active fraction of the population, thereby releasing nutrients that reactivate dormant cells. This in turn may enable a second wave of phage replication and lysis, supporting sustained phage propagation even in the absence of external nutrient input.

These findings suggest that ΦCaCom2 may possess unique ecological or genetic adaptations that allow it to bypass the replication constraints typically faced by phages in stationary-phase bacterial populations. This property enhances its therapeutic potential against *C. acnes* within nutrient-limited niches such as biofilms and sebaceous follicles, where bacterial dormancy is common. Thus, ΦCaCom2 represents a promising candidate for therapeutic interventions targeting antibiotic-resistant or metabolically quiescent *C. acnes* populations.

### Genomic characteristics of the **Φ**CaCom2 phage

The genome of phage ΦCaCom2 was sequenced utilizing Illumina technology, resulting in a single contig of 29,739 base pairs. This contig was determined to be the entire phage genome, with greater than 100-fold coverage (GenBank: OR088869.1). The genome had a single-stranded extension of 13 bases (CCTCGTACGGCTT) at each end, and its overall G+C content was 54.0%, which is lower than that of the *C. acnes* genome (60%). A total of 48 open reading frames (ORFs) were identified. No genes encoding proteins involved in the lysogenic cycle, such as integrases or transcriptional regulators involved in the repression of lysis genes, were discovered. This suggests that ΦCaCom2 phage is incapable of integrating into the host genome and establishing itself as a prophage. *In silico* analysis of the ΦCaCom2 phage genome using PathoFact did not reveal the presence of virulence genes, toxins, or genes encoding proteins involved in antibiotic resistance.

The ΦCaCom2 phage, is taxonomically classified within the recently established Pahexavirus family, under the class Caudoviricetes, according to the latest ICTV taxonomy. Nucleotide similarity analysis using BLASTn revealed that the genomes most closely related to phage ΦCaCom2 are *Propionibacterium* phage Solid (NC_027627.1) with 91.56% identity, *Propionibacterium* phage Cota (MN329678.1) with 91.12% identity, and PA6 (NC_009541.1) with 92.97% identity at the whole-genome level (see Supplementary Fig. S3 online).

### The phage **Φ**CaCom2 is lytic and has the ability to establish a pseudolysogenic cycle in *C. acnes*

Our analysis showed that the ΦCaCom2 phage formed lysis plaques with a turbid center and surrounding transparent halo in 70.8% of the 24 analyzed strains (Fig. 3A and Table 1).

**Figure 3.**
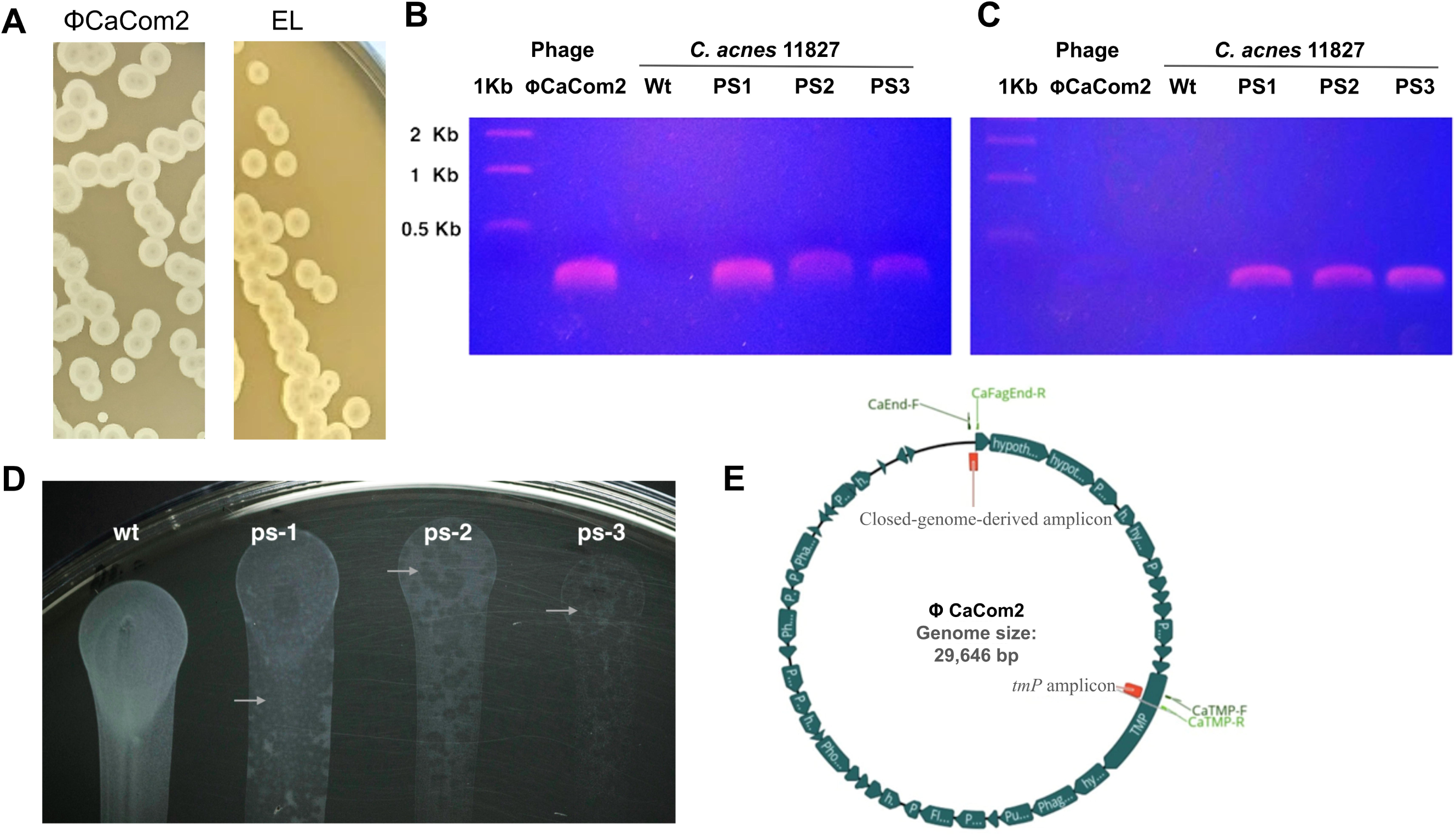
T**h**e **phage** Φ**CaCom2 is lytic and has the ability to establish a pseudo-lysogenic cycle in *C. acnes*. (A**) Morphology of lysis plaques formed on *C. acnes* 11827 by the ΦCaCom2 phage and a phage (El) isolated from a cosmetic product (ElisDay Serum). Note that both phages produce plaques with a turbid center due to the growth of phage-resistant cells. (**B**) Phage-resistant isolates (PS1, PS2, and PS3) were obtained from *C. acnes* 11827 after lytic infection with the ΦCaCom2 phage. The presence of phage DNA in isolates exhibiting superinfection immunity was determined by PCR using primers targeting the phage *TMP* gene. The DNA isolated from purified ΦCaCom2 phages was included as a positive amplification control, while the DNA from *C. acnes* 11827 wild type (Wt) was included as a negative control to demonstrate the specificity of the primers. (**C**) Detection of the circularized genome of the ΦCaCom2 phage in pseudolysogens. Primers were designed to hybridize at each end of the linear ΦCaCom2 genome, generating an amplicon only when the genome is circularized. The DNA isolated from purified ΦCaCom2 phages and DNA from *C. acnes* 11827 wild type (Wt) were included as negative controls to demonstrate the specificity of the primers. (**D**) The image shows that the growth of the pseudolysogens (PS1, PS2, and PS3) is lower compared to the parental strain (Wt) after two days of incubation on FBS agar. The arrows indicate lysis zones in the pseudolysogen lawns. (**E**) The diagram shows the circularized genome map of the ΦCaCom2 phage, indicating the position of the primers and the amplicons (red boxes) obtained for the characterization of pseudolysogeny.

Using *C. acnes* 11827 as the host strain, cells were extracted from the turbid zone of lysis plaques and streaked to cultivate isolated colonies. Subsequently, 30 colonies were randomly selected and re-isolated three times consecutively. Notably, approximately 30% of the colonies exhibited resistance to the ΦCaCom2 phage, while the remaining 70% were sensitive to phage infection. PCR analysis of the phage-resistant colonies confirmed the presence of the phage genome in a circular form (Fig. 3B-C).

Collectively, these results indicate that ΦCaCom2 can establish pseudolysogeny in a subset of *C. acnes* strains. Notably, pseudolysogeny was not observed in 7 out of the 24 strains evaluated (Table 1), in agreement with prior observations reported for *C. acnes* phages. 14. The growth localized in the lysis zone suggests that pseudolysogeny enables the development of a superinfection resistance (SIR) process, which could originate from the superinfection exclusion mechanism (SIE).

Three colonies derived from strain 11827 and harboring the phage genome in a circular form were randomly selected. These pseudolysogens were designated PS1, PS2, and PS3. Interestingly, we observed that these pseudolysogens exhibited slower growth compared to the parental strain when cultured on FBS-agar plates (containing 10% fetal bovine serum and 1.5% agar) (Fig. 3D). Additionally, lysis plaques appeared in agar cultures, suggesting the spontaneous induction of the lytic cycle. This indicates that the pseudolysogenic state is unstable and imposes a fitness cost on the bacterium.

Ellis Day Phage Serum” is a cosmetic product marketed in the USA that utilizes phages to target and eliminate *C. acnes* as a treatment for acne. We observed that phages isolated from phage serum formed turbid lysis plaques with the same morphological characteristics as those formed by the ΦCaCom2 phage (Fig. 3A). These findings are in line with those of previous research suggesting that phages infecting *C. acnes* tend to follow a pseudolysogenic mode of replication and lead to the development of SIE14,16,17.

### The pseudolysogeny process affects biofilm formation in *C. acnes*

Biofilm formation is a well-recognized virulence factor among pathogenic bacterial species, including *C. acnes*33,33–35. In this study, we quantified biofilm formation in three pseudolysogens derivatives of strain 11827 and three derived from strain YJIA, which was isolated from an acne patient. Interestingly, we found that the YJIA strain exhibited greater biofilm biomass compared to the laboratory strain 11827 (Fig. 1B and 4A). Furthermore, the pseudolysogens displayed a reduced capacity to develop biofilms relative to their parental counterparts (see Fig. 4a and Supplementary Fig. S2).

**Figure 4.**
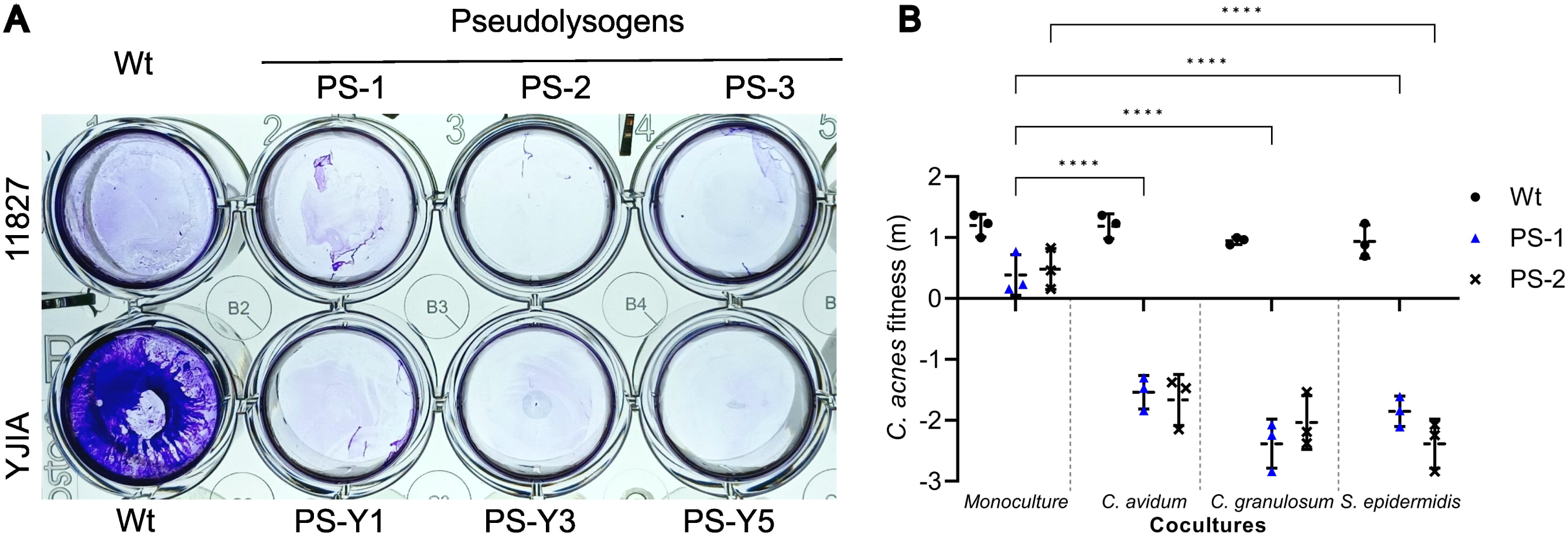
Pseudolysogeny in *C. acnes* alters biofilm formation and ecological fitness against skin commensals. (**A**) Representative crystal violet-stained biofilms formed over 72Lh by ancestral *C. acnes* strains (11827 and JIA) and their corresponding pseudolysogenic (PS) derivatives. Biofilms were developed in polystyrene tissue culture plates under static conditions. **(B)** Comparative fitness (Malthusian parameter, *m*) of the parental strain 11827 (WT) and pseudolysogens (PS-1, PS-2), grown either in monoculture or in co-culture with skin commensal bacteria. Cultures were incubated in heat-inactivated fetal bovine serum, and bacterial growth was quantified by CFU/mL at 72Lh. Data normality was assessed using the Shapiro–Wilk test. Statistical analysis was performed using two-way ANOVA followed by Tukey’s multiple comparisons test, comparing each pseudolysogen in co-culture to its respective monoculture. Asterisks indicate statistically significant differences (*p*L<L0.0001). No significant differences were found between monoculture and co-culture conditions for the wild-type strain. Values represent mean ± standard deviation from three independent experiments (*n*L=L3).

### Pseudolysogeny has a high fitness cost in *C. acnes*

To investigate how bacterial species inhabiting the skin can influence the fitness of pseudolysogens, we conducted paired co-cultures of the parental strain or pseudolysogens of *C. acnes* with *Staphylococcus aureus*, *S. epidermidis*, or *C. avidum* in fetal bovine serum.

Cultures inoculated with pseudolysogens were infected with 2×10^8^ PFU/mL of the ΦCaCom2 phage as a selection pressure to prevent the growth of cells that lost the phage genome. *C. avidum* colonies were differentiated from *C. acnes* by their ability to grow under aerobic conditions and form larger white colonies than *C. acnes* 11827 colonies on BHI-blood agar. Interestingly, we found that the fitness (Maltusian parameter, *m*) of pseudolysogens in the co-cultures was negative and lower compared to the fitness in monocultures. In contrast, the fitness of the parental strain was not affected in the cocultures (Fig. 4B). Additionally, the growth of pseudolysogens in pure cultures was significantly lower than that of the parental strain in fetal serum and solid BHI (Fig. 3D). Overall, these results demonstrate that pseudolysogeny negatively affects the fitness of *C. acnes*, causing pseudolysogens to be non-competitive in cocultures with other bacterial species.

### Pseudolysogeny eliminates clindamycin and erythromycin resistance and does not generate antibiotic-resistant mutants

Macrolides is a widely prescribed treatment for acne, but antibiotic resistance is a growing concern.We found that 9 out of 22 isolates (40.9%) obtained from acne patients are resistant to clindamycin (Supplementary Table S1) and erythromycin. In this study, we evaluated the antibiotic sensitivity of 14 pseudolysogens derived from 7 clindamycin -resistant *C. acnes* strains using the disk diffusion method (Supplementary Table S2). Interestingly, we found that all 14 pseudolysogens had lost their resistance to clindamycin and erythromycin (Fig. 5A and 5B) and remained sensitive to all the other antibiotics tested (Supplementary Table S2).

**Figure 5.**
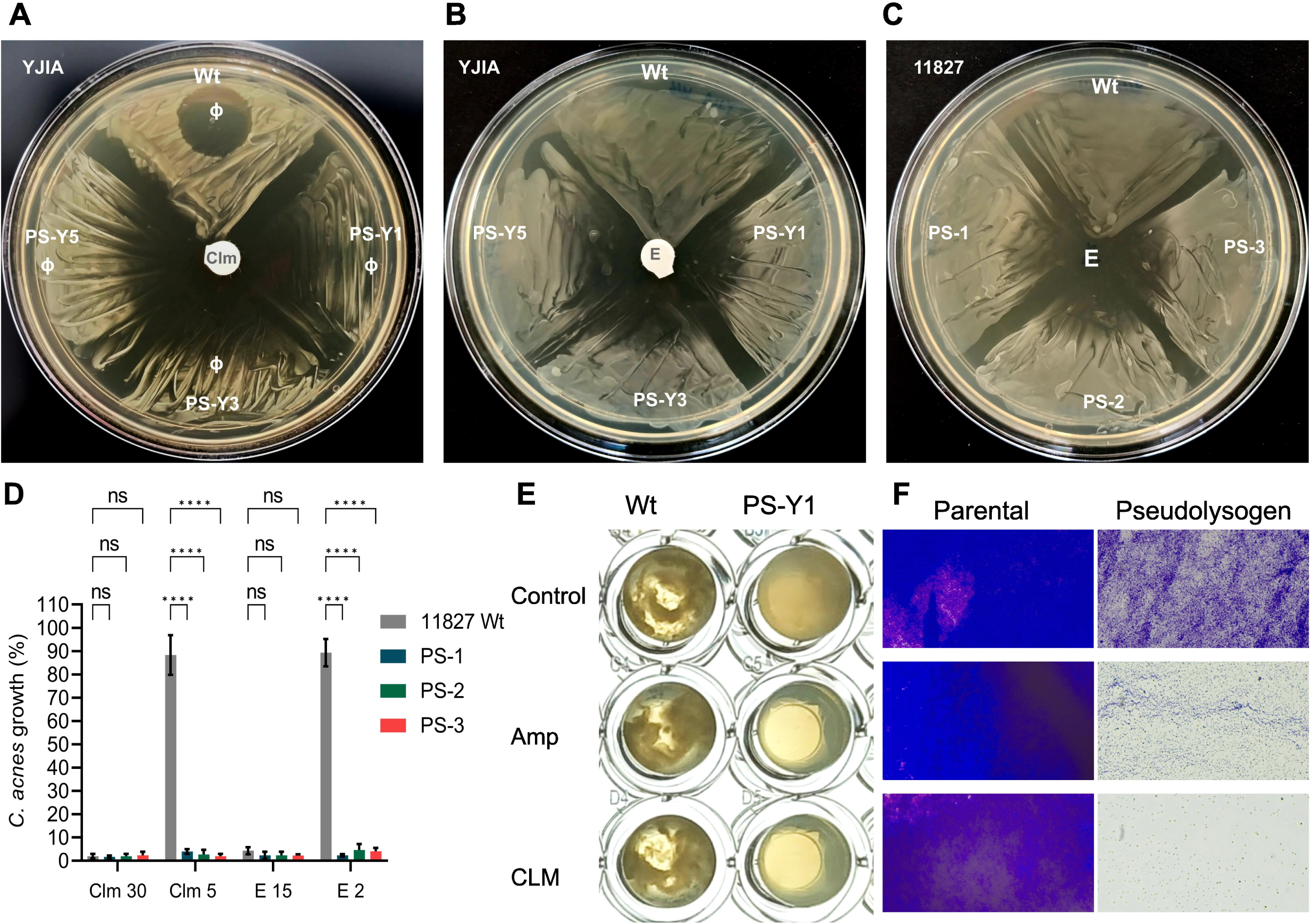
Pseudolysogeny in *C. acnes* reverses resistance to clindamycin and erythromycin. (**A**) Lysis assay showing that ΦCaCom2 phage (Φ) forms clear plaques on lawns of *C. acnes* YIJA wild-type (wt), but not on its pseudolysogenic derivatives (PS-1Y, PS-3Y, PS-Y5), indicating superinfection immunity. Disk diffusion assay shows resistance to clindamycin (30Lµg) in the YIJA wt strain, while pseudolysogens display restored sensitivity. (**B**) Similar results are observed for erythromycin (15Lµg), with pseudolysogens losing the resistance phenotype present in the YIJA wt strain. (**C**) Pseudolysogens derived from *C. acnes* 11827 are sensitive to 1.5Lµg/mL erythromycin, a concentration at which the wild-type strain remains resistant. **(D)** Quantitative assessment of antibiotic susceptibility in liquid culture. Growth of *C. acnes* 11827 wild-type (WT) and pseudolysogens (PS-1, PS-2, PS-3) was measured after 24Lh of exposure to clindamycin (30 or 5Lµg/mL) or erythromycin (15 or 2Lµg/mL). Growth is expressed as a percentage relative to untreated controls. Data are presented as mean ± standard deviation (SD) from three independent experiments (*n*L=L3). Prior to statistical analysis, normality was assessed using the Shapiro–Wilk test (*p* > 0.05). Statistical comparisons were performed using two-way ANOVA followed by Dunnett’s multiple comparisons test. Asterisks indicate statistically significant differences (***p*** < 0.0001) between pseudolysogens and the wild-type strain under the same antibiotic treatment. (**E**) Physical appearance of 72-hour *C. acnes* YIJA-wt cultures grown in polystyrene well plates with BHI broth under static conditions. The wild-type (Wt) strain forms macroscopic cellular aggregates, while the pseudolysogen PS-Y1 exhibits a homogeneous cell distribution. Treatment with 10Lµg/mL ampicillin (Amp) or 30 µg/mLclindamycin (Clm) significantly inhibits pseudolysogen growth but does not affect the wild-type strain. (**F**) Representative 100× micrograph of biofilms formed at the bottom of the wells shown in panel (**E**).

The wild-type *C. acnes* 11827 strain is susceptible to 30 µg/mL clindamycin but exhibits resistance at 2 µg/mL, a subinhibitory concentration. In contrast, pseudolysogens derived from this strain were sensitive even at the subinhibitory concentration (Fig. 5C and 5D). These findings suggest that pseudolysogeny significantly reduces tolerance to sublethal doses of clindamycin.

Furthermore, we found that the YIJA wild-type strain, which was susceptible to ampicillin in BHI agar diffusion assays, exhibited resistance to ampicillin in BHI broth (Fig. 5E and 5F). Conversely, the growth and biofilm formation of the pseudolysogens were sensitive to the inhibitory effects of ampicillin (Fig. 5E and 5F). These findings suggest that the pseudolysogeny process negatively impacts the resistance phenotype and does not seem to promote the emergence of resistant mutants to the antimicrobials evaluated under *in vitro* conditions.

### The **Φ**CaCom2 phage is unable to transduce clindamycin resistance

Genetic transduction is one of many mechanisms of horizontal gene transfer. Several studies have shown that phage-mediated transduction significantly contributes to the dissemination of antibiotic resistance genes36. Here, we evaluated the ability of phage ΦCaCom2 to mobilize clindamycin resistance from strain YIJA to strain 11827. Transduction assays were performed at multiplicities of infection (MOI) of 0.1, 1, and 5, exposing recipient cells to phages for 24 and 48 hours. Clindamycin-resistant cells were not recovered in any of the transduction assays, suggesting that phage ΦCaCom2 either lacks transduction capability or exhibits a very limited ability to transfer clindamycin resistance.

### The phage formulation did not exhibit cytotoxic effects in experiments using human cell cultures

The safety of the ΦCaCom2 phage formulation was evaluated by exposing cultures of human keratinocytes to high (2×10^9^), medium (2×10^8^), and low (2×10^6^) concentrations of the ΦCaCom2 phage for 72 hours, as detailed in the supplemental material and methods. Low and medium concentrations of phages did not significantly affect (p > 0.05) the viability of HaCat cells. Exposure to 2×10^9^ PFU/mL only reduced cell viability by approximately 5% (see Supplementary Fig. S4 online). This shows that the topical formulation with the ΦCaCom2 phage has no cytotoxic activity, even at a high phage concentration.

### Evaluation of the efficacy and safety of **Φ**CaCom2 bacteriophage therapy in patients with mild to moderate acne

We conducted a study to assess the efficacy of bacteriophage ΦCaCom2 in reducing populations of *C. acnes* and to understand the benefits of phage therapy in eighteen mexican patients with mild to moderate acne vulgaris. Of the initially enrolled 20 subjects, 18 (7 males and 11 females) completed the study; two withdrew due to personal reasons. The mean age of the participants was 30 years old. Patients applied approximately 200 µL of MS-C formulation with 2×10^8^ PFU/mL of ΦCaCom2 phage on the entire face, once nightly.

The safety of phage therapy was assessed using local tolerability scoring and adverse event monitoring. For facial acne, the worst score for local tolerability signs and symptoms compared to baseline was observed in 5% (1/18) of patients (erythema: 0%, scaling: 0%, dryness: 5%, and stinging/burning: 0%). Starting from the second week of treatment, a 17.6% reduction in inflammatory lesions and a 15.7% reduction in non-inflammatory lesions were observed.

A statistically significant reduction in both inflammatory and non-inflammatory lesions was observed after 4 weeks (*p*L<0.005). No significant changes were detected during the first and second weeks of treatment. The greatest reduction in inflammatory (−65.74%) and non-inflammatory (−66.32%) lesions occurred after 12 weeks of phage treatment (p<0.0001).These results indicate that the ΦCaCom2 phage formulated in a topical aqueous solution provided significant benefits as early as the second week of treatment (Fig. 6A and 6B).

**Figure 6.**
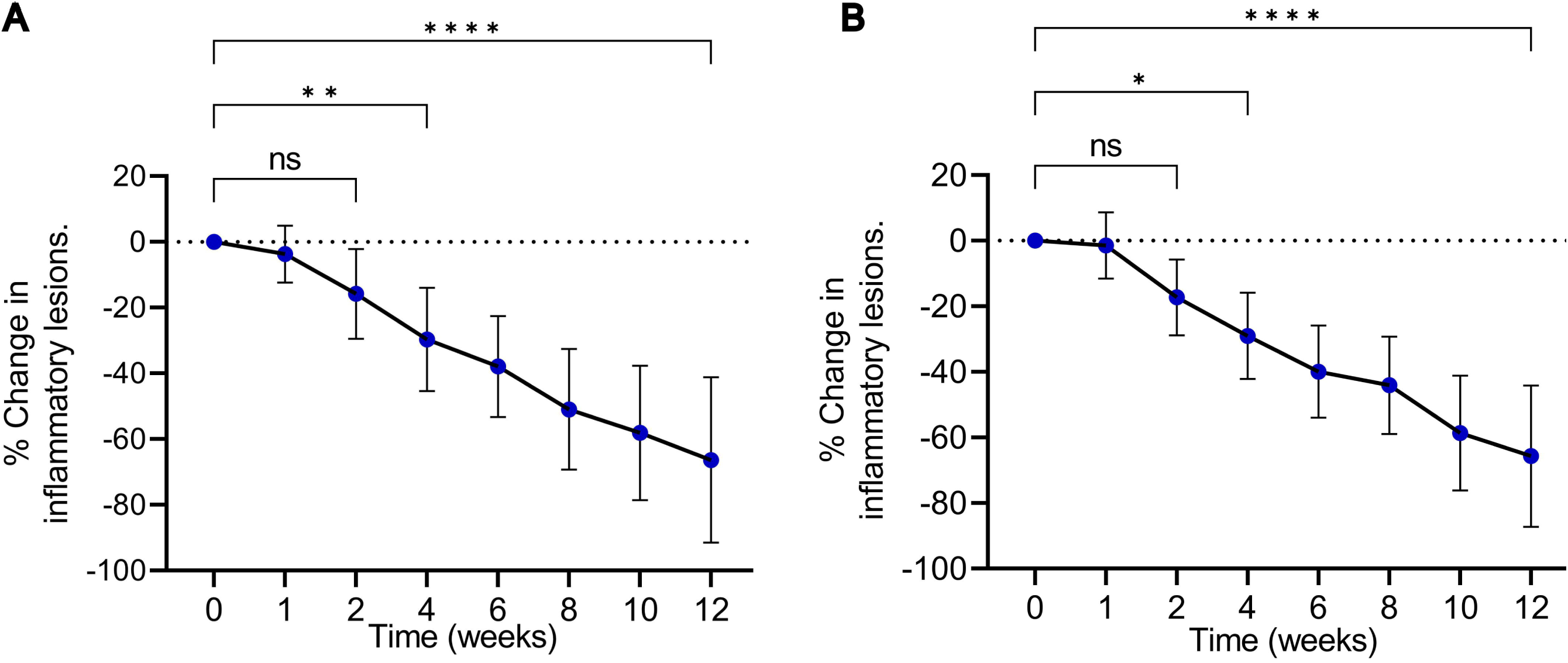
Quantitative reduction of facial acne lesions following topical phage therapy. Topical application of a ΦCaCom2 phage-based formulation significantly reduced (**A**) inflammatory and (**B**) non-inflammatory facial acne lesions in volunteers with mild to moderate acne. The graphs show the percentage change in lesion count relative to baseline (day 0), with baseline values normalized to 100%. Negative values indicate a reduction in lesion count compared to baseline. Statistical significance was assessed using the Friedman test for repeated measures. Post hoc comparisons were performed using Dunn’s test with correction for multiple comparisons (αL=L0.05). Percentage change at each time point was compared to baseline. A statistically significant reduction in both inflammatory and non-inflammatory lesions was observed after 4 weeks (\**p*L<L0.05, **p<0.001, **** *p*L<L0.0001). No significant (ns) changes were detected during the first and second weeks of treatment. Data are presented as mean ± standard deviation (n = 18 volunteers).

After two months of topical application of phage ΦCaCom2, volunteers showed marked clinical improvement. This was supported by photographic documentation of facial acne lesions taken before and after treatment (Fig. 7), which revealed consistent reductions in lesion severity and overall improvement in skin condition across individuals.

**Figure 7:**
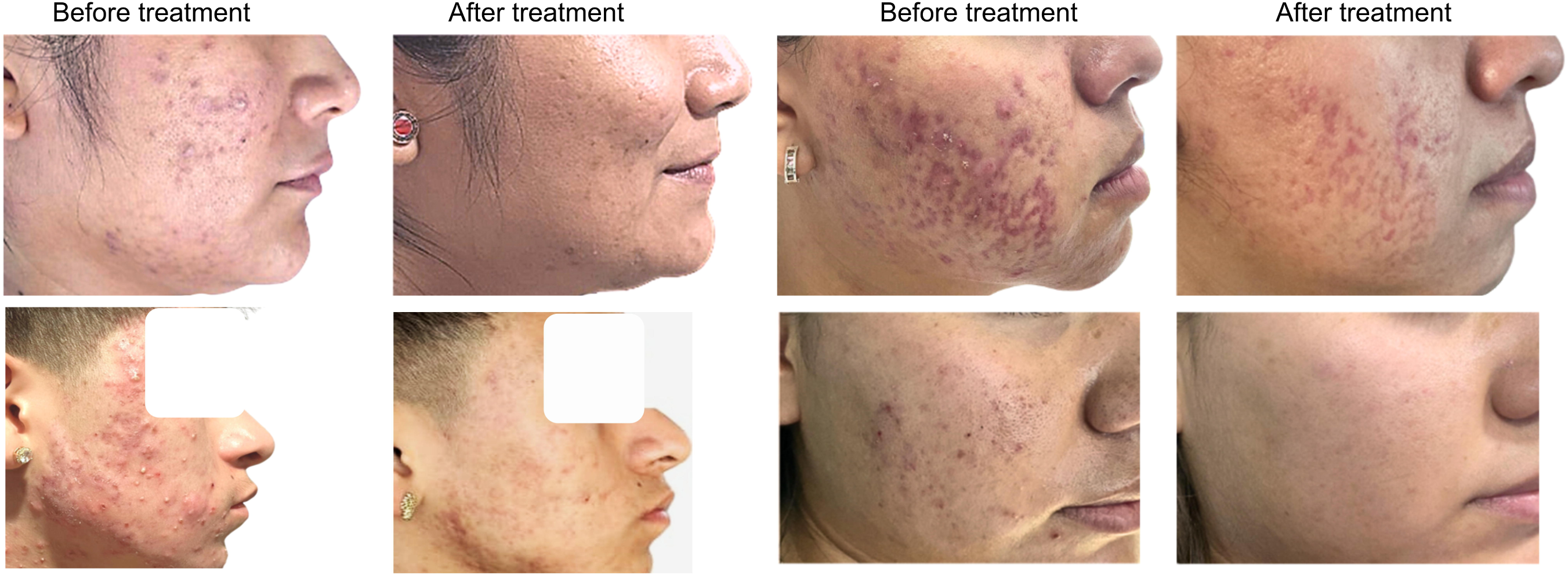
Clinical improvement in acne following topical phage treatment. Representative clinical images of individuals with varying acne severity before and after two months of topical application of phage ΦCaCom2. Treatment resulted in visible reductions in inflammatory lesions, erythema, and overall skin irritation.

### Topical application of the **Φ**CaCom2 phage reduces the abundance of *C. acnes* on the skin

The efficacy of ΦCaCom2 phage in reducing the abundance of *C. acnes* on the facial skin was evaluated in 18 volunteers by estimating CFU/cm². Samples were collected one day before treatment initiation (baseline), at the end of the 12-week treatment period, and two weeks post-treatment. Baseline abundance was normalized to 100% for each participant. Phage therapy resulted in a significant reduction in *C. acnes* levels, with an average decrease of 31% compared to baseline (p < 0.0001). Notably, two weeks after treatment completion, *C. acnes* abundance remained significantly lower than baseline levels (p<0.0001; Fig. 8A).

**Figure 8.**
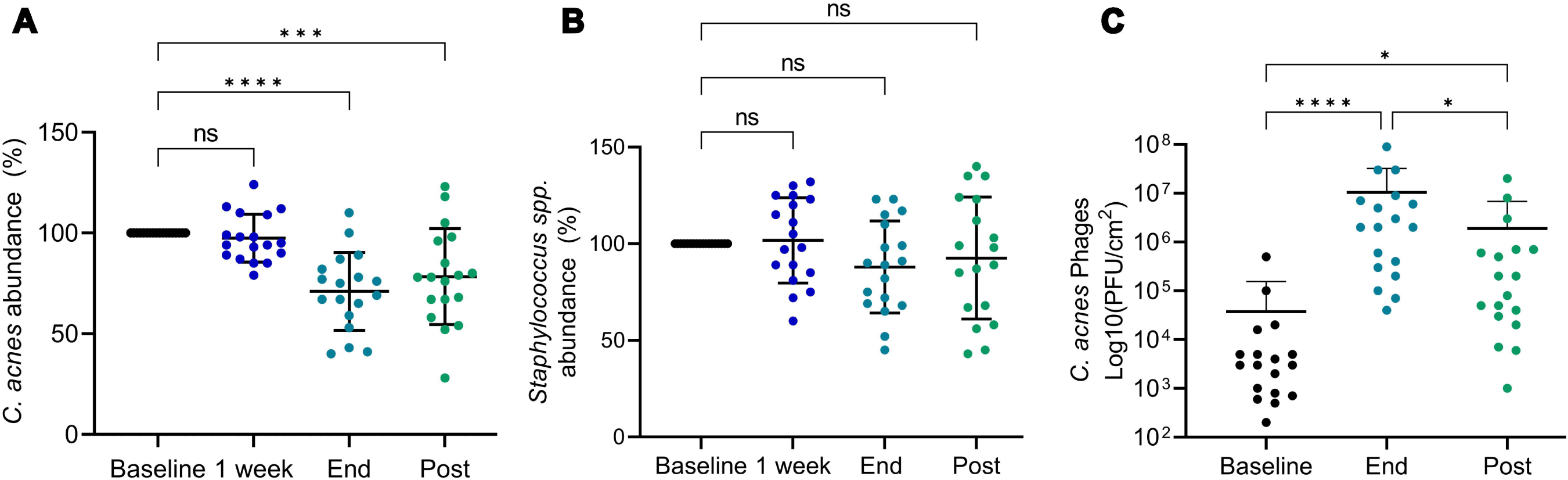
Topical application of. Φ**CaCom2 phage selectively reduces *C. acnes* populations without altering *Staphylococcus spp.* abundance.** (**A**) The mean percentage reduction in viable *C. acnes* populations was assessed at three time points: one week after treatment initiation (1 week), at treatment completion (End), and two weeks post-treatment (Post). The baseline value (100%) is shown for reference. For each volunteer, *C. acnes* colony-forming unit (CFU/cm²) density measured one day before treatment was normalized to 100%. P-values obtained using the Friedman test followed by Dunn’s multiple comparisons test are indicated as follows: \*\**p* = 0.0009 and \*\*\**p* < 0.0001, relative to baseline. No statistically significant differences (ns) were observed between baseline and the 1-week time point. Data are presented as mean ± standard deviation (n = 18 volunteers). (**B**) The mean percentage abundance of viable *Staphylococcus* spp. on the skin was quantified one day before and at the end of the phage treatment period to assess off-target effects. *Staphylococcus* spp. CFU/cm² density measured before treatment was normalized to 100% and is shown as the baseline reference. Data are presented as mean ± standard deviation (n = 18 volunteers). No statistically significant differences were found between time points (p > 0.05), as determined by the Friedman test followed by Dunn’s multiple comparisons test. (**C**) The density of phages infecting *C. acnes* 11827 (PFU/cm²) was quantified at baseline (before treatment), at the end of treatment (End), and four weeks after treatment completion (Post). P-values obtained through the Friedman test followed by Dunn’s multiple comparisons test are indicated as follows: *p* < 0.05 and \*\*\**p* < 0.0001, relative to baseline. Data are presented as mean ± standard deviation (n = 18 volunteers).

The sustained reduction in *C. acnes* population observed two weeks after treatment suggests that phages are capable of colonizing the skin and persisting in situ, thereby controlling *C. acnes* overgrowth. This effect is consistent with the significant decrease in both inflammatory and non-inflammatory lesions observed in the treated volunteers.

### No phage-resistant cells were detected in patients at the end of the treatment

We investigated the potential emergence of phage-resistant *C. acnes* variants during the phage treatment. At week 12 of phage treatment, a total of 540 *C. acnes* colonies (30 colonies per volunteer) were isolated to quantify the emergence of cells with phage resistance mechanisms. Notably, all *C. acnes* colonies identified by PCR were found to be sensitive to the ΦCaCom2 phage. These results suggest that populations of pseudolysogens with a superinfection resistance phenotype were absent or undetectable *in vivo*. We hypothesize that the absence of pseudolysogens *in vivo* is due to the fitness cost associated with pseudolysogeny-mediated resistance, which prevents the establishment of a population capable of competing with wild-type cells, the skin microbiome species, and the pressures imposed by the human host’s antimicrobial mechanisms.

### The reduction of *C. acnes* populations by phages did not lead to an increase in *Staphylococcus* species abundance on the skin surface

Evidence suggests that *C. acnes* competes with *Staphylococcus* spp. and can suppress their abundance on human skin<u>37</u>. To determine whether phage-mediated reduction of *C. acnes* populations alters *Staphylococcus* levels, we quantified *Staphylococcus* spp. CFU/cm² one day before treatment, one week after treatment initiation, at treatment completion, and two weeks post-treatment. Despite a significant decrease in *C. acnes*, no increase in *Staphylococcus* spp. abundance was observed on the skin surface or within acne lesions following ΦCaCom2 phage application (Fig. 8B).

### Abundance of phages targeting *C. acnes* increases and persists on the skin of acne-affected volunteers following treatment

To evaluate phage persistence and changes in their abundance, skin samples were collected at baseline, at the end of treatment, and two weeks post-treatment. The results showed a marked increase in phages infecting *C. acnes* 11827 during the treatment period, with elevated levels maintained after its completion (Fig. 8C). These findings indicate that the applied phages replicate effectively on the skin and remain viable despite routine cleansing, supporting their potential for sustained therapeutic action.

## Discussion

Our study demonstrates that pseudolysogenic phages targeting *C. acnes* can reduce bacterial burden while imposing substantial fitness costs that constrain the emergence of phage resistance. These findings challenge the prevailing notion that only strictly lytic phages are suitable for therapeutic applications and highlight pseudolysogeny as a biologically advantageous mechanism to limit *C. acnes* virulence and antibiotic resistance *in vivo*.

A central outcome of this work is the identification of evolutionary trade-offs mediated by pseudolysogeny, wherein *C. acnes* cells that adopt a pseudolysogenic state to evade lysis exhibit diminished biofilm formation and compromised competitiveness within the skin microbiome. Biofilm formation is a critical determinant of *C. acnes* persistence and antibiotic tolerance 34,35; thus, its attenuation following pseudolysogenic infection suggests a dual benefit of this phage strategy: direct bacterial reduction and long-term suppression of virulent phenotypes. This is consistent with theoretical frameworks and empirical evidence in other bacterial systems, where resistance to phage predation is often accompanied by fitness penalties38–40.

Importantly, unlike observations from other phage therapy models where bacterial resistance rapidly emerges and compromises efficacy 41, no phage-resistant *C. acnes* isolates were detected after treatment. This likely reflects the high fitness cost of pseudolysogeny and SIE, which, while providing temporary superinfection resistance, severely limits bacterial capacity to thrive in the competitive skin environment. These results align with studies in other microbiomes showing that phage-mediated selection can promote trade-offs that favor susceptible, rather than resistant, bacterial populations over time40 .

Our findings also have important implications for microbiome stability and the safety of phage therapy. Despite the marked reduction in *C. acnes* populations, the abundance of *Staphylococcus* spp.—key commensals involved in skin homeostasis42——remained unaffected. This supports the notion that *C. acnes* phages exhibit high specificity with minimal collateral disruption, offering a key advantage over broad-spectrum antibiotics43 11. However, while we assessed *Staphylococcus* spp. to evaluate off-target effects, a key limitation of our study is the exclusion of non-cultivable microorganisms. Broader microbiome shifts may have gone undetected, underscoring the need for future metagenomic analyses to comprehensively assess the impact of phage therapy on skin microbial communities.

Transduction assays revealed that phage ΦCaCom2 does not mediate horizontal transfer of clindamycin resistance under the tested conditions. This supports its suitability for therapeutic use, as it minimizes the risk of spreading antibiotic resistance genes. However, since only clindamycin resistance was evaluated and under specific in vitro conditions, the possibility of low-frequency or context-dependent transduction events cannot be entirely ruled out. Further studies under physiologically relevant conditions are needed to confirm the absence of transduction capacity.

The persistence of viable phages in the skin microbiome for at least four weeks post-treatment raises the possibility of long-term ecological effects, wherein phages may act as sustained regulators of *C. acnes* populations. While persistent pseudolysogenic phages could theoretically interfere with subsequent therapeutic interventions, the observed reduction in virulence-associated traits suggests that such persistence may be beneficial. This contrasts with concerns in other systems where lysogeny-associated immunity can limit phage therapy success44,45.

Interestingly, while pseudolysogeny was stable *in vitro*, we did not recover pseudolysogens from clinical samples. This may be due to asymmetric segregation of the phage genome during cell division46 and the ecological disadvantages associated with pseudolysogeny, which likely prevent stable maintenance *in vivo*. We propose that pseudolysogeny functions as an eco-evolutionary constraint: it offers transient advantages like superinfection resistance 17, while imposing costs such as impaired biofilm formation, reduced interspecies competitiveness, and increased antibiotic susceptibility.

Although our study focused on a single pseudolysogenic phage, the broader relevance of these findings warrants further investigation. Given the widespread pseudolysogenic nature of *C. acnes* phages [4,6], it is plausible that this interaction confers selective advantages under specific environmental or physiological conditions46,47. Future studies should aim to elucidate the molecular mechanisms underlying these fitness trade-offs and assess how pseudolysogeny shapes interactions with other skin microbes, microbial resilience, and host responses.

In conclusion, our findings establish pseudolysogeny as a key eco-evolutionary constraint that enhances the efficacy and safety of phage therapy for acne vulgaris. By combining direct bacterial suppression with evolutionary trade-offs that limit resistance and virulence, pseudolysogenic phages represent a promising strategy for microbiome-informed therapeutics. Future research should focus on optimizing phage therapies that harness these dynamics and assessing their long-term ecological effects. Moreover, investigating pseudolysogeny as a regulatory mechanism in natural microbiomes may reveal fundamental principles governing microbial community structure, stability, and evolution.

## Acknowledgements.

We are grateful to all the volunteers for their participation in the studies. Our sincere appreciation goes to dermatologist Dr. Gissel Castellanos Ramos for their assistance in setting up the clinical study. We also thank Andres Andrade-Trejo for his support in collecting the samples used for phage isolation. Additionally, we appreciate the contributions of Jessica G. Aguilar and Emiliano Román to the isolation and characterization of *C. acnes* strains. Special thanks to Raúl Quijada Ibarra for his help in annotating and analyzing the CaCom2 phage genome.

## Author contributions

A.T. and A.A. conceptualized and designed the study, performed data analysis, and drafted the manuscript. A.T. also contributed to data collection, statistical analysis, and critical revision of the manuscript. A.C. assisted in the interpretation of results, manuscript editing, and overall project supervision. P.S. contributed to the study design and conducted the clinical outcome assessments. All authors reviewed and approved the final version of the manuscript.

## Data availability

The ΦCaCom2 genome sequence and annotation are deposited in the NCBI Genbank database under accession number OR088869.1. Additionally, the partial sequence of the 16S ribosomal RNA gene is deposited in the NCBI Genbank under the accession numbers listed in Table 2.

**Table 2.**
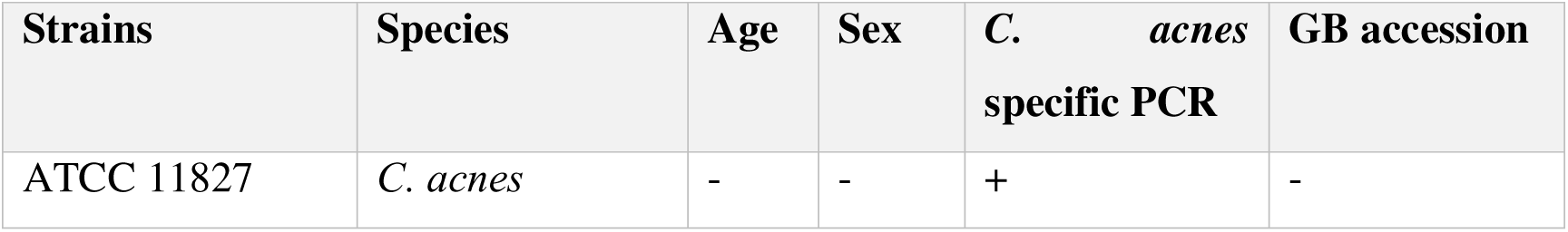
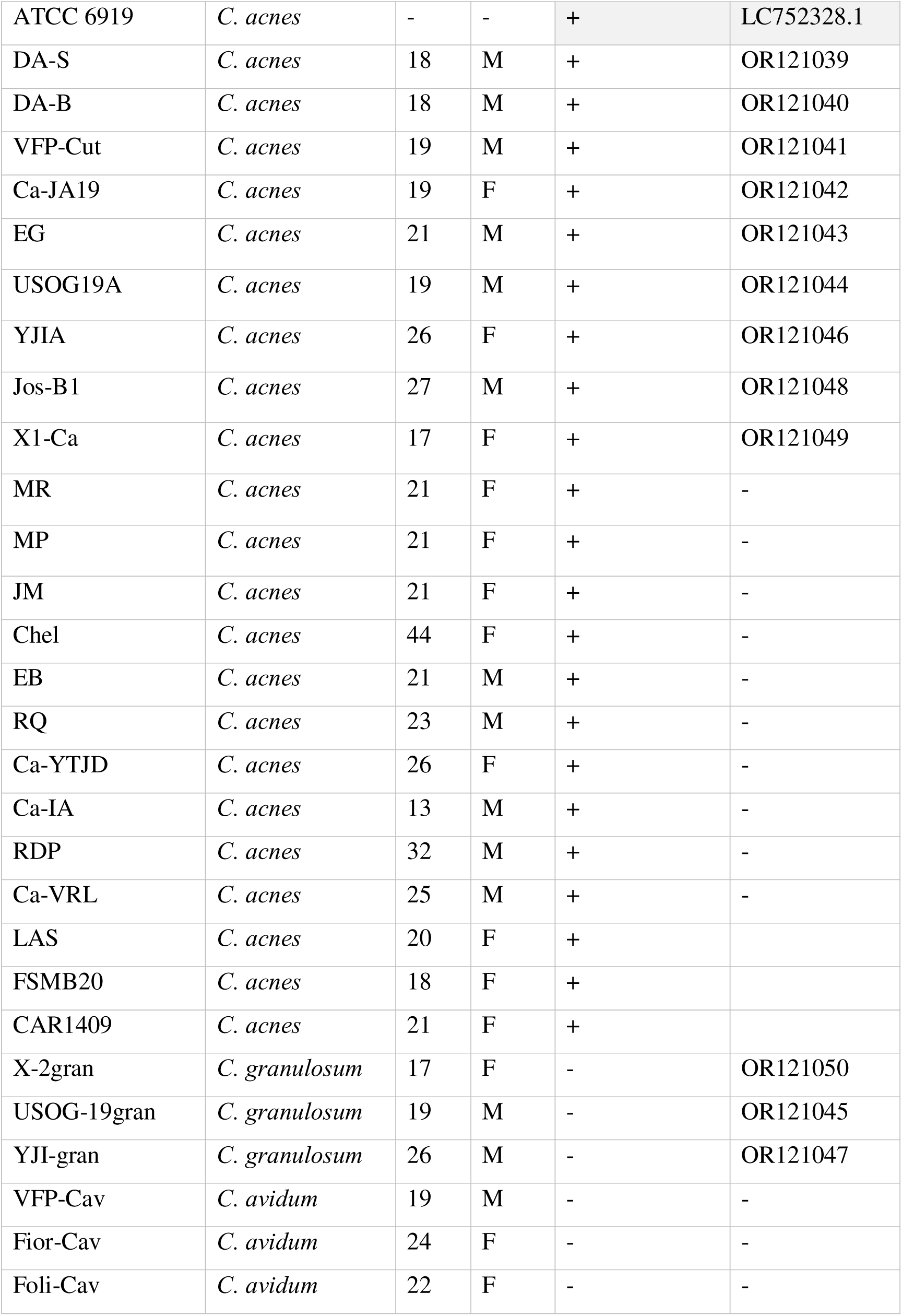
Origin and characteristics of *Cutibacterium* strains used as reference and isolates from acne lesions. *Cutibacterium* species were identified using specific primers for *C. acnes*. The partial sequence of the 16S ribosomal RNA gene was obtained with the MinION sequencer from Nanopore, and the GenBank accession number is provided along with the sex and age of the acne volunteers from whom the strains were isolated.

