## Supplementary Material and Methods for "Pseudolysogeny-mediated evolutionary trade-offs favor phage therapy by limiting antibiotic resistance and virulence in *Cutibacterium acnes*"

**Table of Contents**

Supplementary Material and Methods……………………………….……2

Supplementary Figure S1…………………………………………….9

Supplementary Figure S2…………………………………………….9

Supplementary Figure S3…………………………………………….10

Supplementary Figure S4…………………………………………….10

Supplementary Table S1…………………………………………….11

Supplementary Table S2…………………………………………….12

### Supplementary Material and Methods

#### Identification of *C. acnes* by PCR

The identity of the isolates was determined using specific primers to amplify a region of the 16S rDNA gene of *C. acnes*, resulting in a 600 bp product[1]. The PCR conditions were as follows: one cycle of denaturation at 95 °C for 1 min; 30 cycles of 95 °C for 20 s, 61 °C for 30 s, and 72 °C for 50 s; and one cycle of extension at 72 °C for 1 min.

The reaction mixture consisted of 7.5 µL of 2x Thermo Scientific™ K0171 PCR Master Mix, 1 µL of each oligonucleotide Pa-1 5'-GGCACACCCATCTCTGAGCAC-3' and Pa-2 5'-GGGTTGTAAACCGCTTTCGCTG-3' (10 µM), 1 µL of the lysate of cryopreserved cells, and 4.5 µL of water, resulting in a final volume of 15 µL.

The amplification products were visualized on a 1% agarose gel. The running buffer used was TBE, and 15 µL of the reaction was mixed with 2 µL of 10x DNA loading buffer containing the GelRed fluorophore. Electrophoresis was conducted at 100 volts for 30 minutes. The gel was exposed to ultraviolet light in a transilluminator and photographed using a smartphone camera.

#### Nanopore sequencing of the 16S rRNA gene for bacterial identification.

DNA was extracted from the samples using the DNeasy PowerSoil Kit (QIAGEN, # 12888-100) following the manufacturer's instructions. The 16S ribosomal RNA (rRNA) gene was amplified by PCR using the 16S barcoding kit 1-24 (SQK-16S024) as described in the ONT protocol, 25 µL of LongAmp hot start Taq 2× master mix (New England Biolabs, MA, USA) was mixed with 14 µL of nuclease-free water, 1 µL of DNA from pure isolates, and 10 µL of a 16S barcode. These barcodes use the 27F (5′-AGAGTTTGATCMTGGCTCAG-3′) and 1492R (5′-CGGTTACCTTGTTACGACTT-3′) primers to amplify the full-length 16S rRNA bacterial gene. PCR was then conducted on a Eppendorf thermal cycler (Mastercycler Pro 6321, Hamburg, Germany) using the following conditions: 1 cycle of 95°C for 1 min, 33 cycles of 95°C for 20 s, 55°C for 30 s, and 65°C for 2 min, and a final extension of 65°C for 5 min. Three microliters of the reaction were analyzed by agarose gel electrophoresis to confirm the presence of an amplicon of approximately 1.5 Kb in size. The PCR products were purified using the High pure PCR product Purification kit (Ref 11732676001) from ROCHE. The DNA concentration and purity were determined using an EPOCH spectrophotometer.

Library preparation for bacterial 16S rRNA gene sequencing on the MinION Mk1c (MC-112236) portable sequencing device (Oxford Nanopore Technologies, Oxford, UK) was conducted using the 16S barcoding kit 1-24 (SQK-16S024) and the "rapid sequencing amplicons—16S barcoding protocol" version 16S_9086_v1_revU_14Aug2019 with some minor modifications.

The libraries were sequenced with a FLO-MIN106 flow cell. Sequencing was initiated through MinKNOW software version 22.10.7 with fast base-calling and a Q score of ≥ 8 for between 5 and 25 h. Once sequencing was stopped, FAST5 reads were base called using the super-high-accuracy base-calling model with barcode removal using Guppy version 6.3.9. To identify the species, each dataset of nanopore reads was analyzed by the EPI2ME (Oxford Nanopore Technologies, Oxford, UK) analysis workflow.

De-multiplexing, adapter trimming, and assembly were performed using Geneious Prime software. The resulting consensus sequence was subsequently subjected to a BLAST search against the GenBank database of 16S rRNA gene sequences. The sequences of each strain were deposited in GenBank (Table 2).

**Antibiotic Susceptibility Testing.**

The antibiotic susceptibility of the isolates was assessed using the disc diffusion assay, following Clinical and Laboratory Standards Institute (CLSI) guidelines with minor modifications. Briefly, five colonies were picked from blood agar plates and suspended in 5 mL of Mueller-Hinton Broth (MHB) (Himedia, India). A sterile swab was immersed in the suspension and used to uniformly inoculate the surface of Mueller-Hinton Agar (MHA) plates (10 cm diameter). A single antibiotic disc was placed on each plate. The antibiotic discs used were: 10 μg ampicillin (AM), 30 μg tetracycline (TE), 30 μg minocycline (MH), 30 μg doxycycline (DO), 15 μg erythromycin (E), and 30 μg clindamycin (CLM) (Oxoid, Germany). Plates were incubated anaerobically at 37 °C for 3 days. After incubation, the diameters of the inhibition zones were measured in millimeters. All experiments were performed in triplicate. Susceptibility was interpreted as resistant or sensitive based on the CLSI 2015 zone diameter breakpoints (Supplementary Table S1).

**Detection of Clindamycin Resistance Genes.**

Detection of *C. acnes* isolates carrying clindamycin resistance genes was performed by PCR amplification of erm(X) and erm(50) genes. The primers used for erm(X) detection were EmRX-F (5′-CTCACCAACCACAAGATCATC-3′) and EmRX-R

(5′-GAAGAGATCGATCCAGTCGTT-3′), and for **erm(50)** detection were erm(50)-F (5′-TCAATGAGGCGACGATCAGAC-3′) and erm(50)-R (5′-GGTGAACACGTCATGGACGA-3′), as previously described [2]. PCR reactions were performed using Thermo Scientific PCR Master Mix 2X (Catalog number K0171) and analyzed by agarose gel electrophoresis.

**Mutations associated with resistance to macrolide.**

23S amplicons were sequenced across the peptidyl transferase region using the internal primer 50 - GTAGCGAAATTCCTTGTCGG-30 [3, 4] .

**Assessment of Phage Host Range Against *C. acnes, C. granulosum* and *C. avidum*.**

The host range of the isolated phages was determined by evaluating their ability to form lysis plaques in double-layer agar assays using BHI medium. Phages were tested against two reference strains of *Cutibacterium acnes* (ATCC 11827 and ATCC 6919), 22 clinical isolates of *C. acnes*, and 3 isolates each of *C. granulosum* and *C. avidum*, all obtained from skin swabs of volunteers with moderate acne.

**Phage amplification.**

The CaCom2 phage was replicated by infecting 500 ml of an exponentially growing culture (~2e8 CFU/ml) of *C. acnes* 11827 strain in BHI liquid medium with ~2e6 plaque forming units (PFU/ml). The culture was incubated under agitation at 150 rpm for 18 hours at 37°C in an anaerobic atmosphere. Non-lysed cells were removed by centrifugation at 8000 rpm for 10 minutes, and the supernatant was filtered using a PVDF membrane with 0.2 µm diameter pores.

**Phage purification for therapeutic use**

A 500 mL lysate of the CaCom2 phage ($7 \times 10^{10}$ PFU/mL), amplified in *C. acnes* 11827, was supplemented with $MgSO_4$ to a final concentration of 5 mM and treated with 100 mg of DNase I (Ref. 11284932001, F. Hoffmann-La Roche Ltd., Basel, Switzerland). The mixture was incubated at 32 °C for 2 hours to degrade residual bacterial DNA. The lysate was subsequently concentrated using an Amicon stirred cell (UFSC400SL, Millipore) fitted with a 100 kDa Biomax polyethersulfone ultrafiltration membrane.

Phages retained on the membrane were first washed with 100 mL of PB-T buffer (distilled water, 150 mM NaCl, 40 mM Tris, pH 7.5, 10 mM $MgSO_4$, 1% Tween 20), followed by an additional wash with 400 mL of PB buffer (identical composition without Tween) to remove residual detergent. Phages were then eluted from the membrane with 100 mL of PB buffer under agitation (50 rpm) for 2 hours at 4 °C.

The final phage suspension was sterilized by filtration through 0.22 µm pore-size polyethersulfone membranes and stored in amber glass vials at 4 °C. Phage titers were determined by serial dilution and plaque assay using *C. acnes* 11827 as the host strain.

Sterility was assessed in triplicate by inoculating 1 mL of the phage suspension into 100 mL of BHI supplemented with 5% defibrinated sheep blood. Three cultures were incubated aerobically and three anaerobically at 37 °C for 14 days. Cultures were inspected every 5 days for evidence of bacterial or fungal growth.

**Thermal Stability of Phages.**

For stability assays, 30 ml of MS-C formulation (Phage buffer ph 7.5 supplemented with polyols and proteins) were inoculated with approximately 6 x 10^8 PFU/mL of phages and stored in amber glass bottles. These bottles were then stored at 4°C in a refrigerator and at 30°C in an incubator for 12 months. To simulate transportation and mishandling conditions, stability was also assessed at 37°C and 42°C for 24 hours. Samples were collected to quantify the viable phage titer through serial dilutions, using *C. acnes* 11827 as the host strain. For each evaluated temperature, three bottles were used, and titrations were performed in triplicate for each sample.

**Static Biofilm Assays.**

The ability for biofilm formation in both the wild type and pseudolysogenic strains was assessed using polystyrene 96-well plates (COSTAR® 10218022). A volume of 1 milliliter of fetal bovine serum was inoculated with 1x107 CFU of cells in exponential phase, with 100 µl added to each well. The same procedure was replicated using BHI medium supplemented with 25 µg/ml of ampicillin. Both sets of plates were then incubated at 37°C for 72 hours.Following the 72-hour incubation period, the BHI medium was meticulously removed to avoid disrupting the biofilm structure. Subsequently, 200 µl of 70% ethanol was introduced to each well to fix the biofilm, undergoing incubation for 5 minutes. After this, the ethanol was extracted, and 200 µl of distilled water was gently applied to rinse off any excess ethanol. Once rinsed, the plates were allowed to air dry within a Biosafety Level II cabinet for 20 minutes.

For the staining process, 100 µl of a 10% crystal violet solution (SIGMA® Cat. #V5265-500ML) was added to each well and allowed to sit for 10 minutes. Following this period, the crystal violet solution was removed, and any excess stain was removed through a series of three washes, each utilizing 200 µl of distilled water.The stained crystal violet was dissolved by adding 150 µL of 95% ethanol, and the optical density (OD) was measured at 595 nm.

**Sampling of the cultivable microbiome.**

Microbiota sampling of specific facial regions was consistently performed by the same investigator during all study visits. The volunteers visited the laboratory for sample extraction, which was promptly processed. To ensure consistency, each sampled area, measuring 1 $cm^2$, was identified using a positioning mask and standardized photography, ensuring the same area was sampled at each follow-up visit. Skin microbiota samples were collected from two areas on the face, including one with comedones and another with papulo-pustular lesions. Sterile cotton-tipped swabs (COPAN SPA, Brescia, Italy) were moistened with PBS and 0.1% Tween 20. The chosen areas were vigorously swabbed for 30 seconds. The cotton swab tip was cut and immersed in 500 µl of BHI medium within a 1.5 ml tube, which was agitated for 1 minute using a vortex. To quantify the cultivable microbiome, 100 µl of the sample was plated on a BHI agar plate supplemented with 5% sheep blood. The plates were incubated under anaerobic conditions at 37°C for 8 days, with reviews conducted every 2 days to record and quantify colony growth. To quantify *Staphylococcus* species populations, 100 µL of the sample was inoculated onto Mannitol salt agar (MSA) (Sigma, ref M9052) plates and incubated for 72 hours at 37 °C.Three representative colonies of each morphology were isolated for identification using selective culture media and biochemical tests.

**Phage Cytotoxicity Assays.**

HaCat cells were seeded at 80 µL per well ($1 \times 10^4$ cells) in 96-well flat-bottom microplates (Costar, Ref. 3603) using RPMI-1640 medium (Ref. 11875-093) supplemented with 10% (v/v) fetal bovine serum (Sigma-Aldrich, Ref. F4135). Cell viability was assessed after 72 hours of incubation under the following conditions: (a) negative control, with no treatment; (b) addition of 20 µL of PB7L formulation; (c) 20 µL of PB7L containing approximately $1\times10^6$ phages; (d) 20 µL of PB7L containing approximately $1\times10^8$ phages; and (e) 20 µL of PB7L containing approximately $1\times10^9$ phages.

Cytotoxicity was evaluated using the 3-(4,5-dimethylthiazol-2-yl)-2,5-diphenyltetrazolium bromide (MTT) assay. This method assesses cell viability, proliferation, and cytotoxicity by quantifying the conversion of the yellow tetrazolium salt into purple formazan crystals by metabolically active cells. The Cell Proliferation Kit I (MTT) (Roche, Ref. 11465007001) was used according to the manufacturer's instructions.

**Eligibility Criteria.**

Both male and female individuals aged 18 years and older were eligible if they had mild to moderate facial acne, as defined by an Investigator Global Assessment (IGA) score of 3 or 4, and lesion counts ranging from 20 to 50 inflammatory lesions (papules, pustules, and nodules), 25 to 100 non-inflammatory lesions (open and closed comedones), and no more than 10 nodules. Pregnant, breastfeeding, or planning to become pregnant women were excluded from the study. The use of oral retinoids or corticosteroids within 12 weeks prior to randomization, as well as the use of topical retinoids, corticosteroids, topical anti-inflammatory agents, oral antibiotics, or other systemic acne treatments within 4 weeks prior to treatment initiation, was prohibited. The use of medicated facial cleansers, topical acne medications, creams, sunscreens, or makeup during treatment was not allowed.

Bacterial culture was conducted to identify only those patients whose lesions were colonized with *C. acnes*. Subjects with lesions harboring more than 30 colony-forming units per lesion of any bacterial species other than C. acnes were excluded.

**Efficacy and Safety Endpoints.**

The co-primary endpoints at Week 8 were the absolute change in counts of inflammatory lesions from baseline and treatment success rates assessed by IGA (Investigator's Global Assessment). The IGA scale was defined as follows: 0 (Normal, clear skin with no evidence of acne vulgaris), 1 (Almost clear, occasional non-inflammatory lesions may be present, with rare non-inflamed papules, resolving papules may be hyperpigmented but not pink-red), and 2 (Mild, some non-inflammatory lesions are present, few inflammatory lesions, papules/pustules only, no noduloquistic lesions).

Safety assessments included treatment-emergent adverse events (TEAE), vital signs, physical examinations, and site tolerability. A subject satisfaction questionnaire was administered at the end of treatment.

**Subjective Patient-reported Efficacy Assessment.**

Subjective self-assessments of acne severity and treatment satisfaction by patients were performed using a visual analog scale (VAS) ranging from 0 (condition at initial visit) to 10 (disease-free state). Treatment-associated irritation was also evaluated using a numerical rating scale ranging from 0 to 10 (0: no discomfort, 10: worst imaginable discomfort).

**Supplementary Figures.**

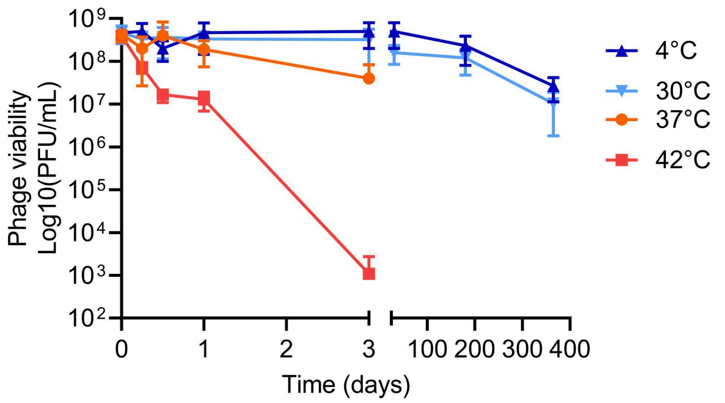


**Supplementary Figure S1. Thermal stability of phages ΦCaCom2 in a topical formulation.** Phage titers were monitored over 12 months at room temperature (24–33 °C) to assess long-term stability. Stability under simulated transport conditions was also tested by incubating phage formulations at 37 °C and 42 °C for 12 hours. The data are presented as mean ± standard deviation (n=3 independent experiments).

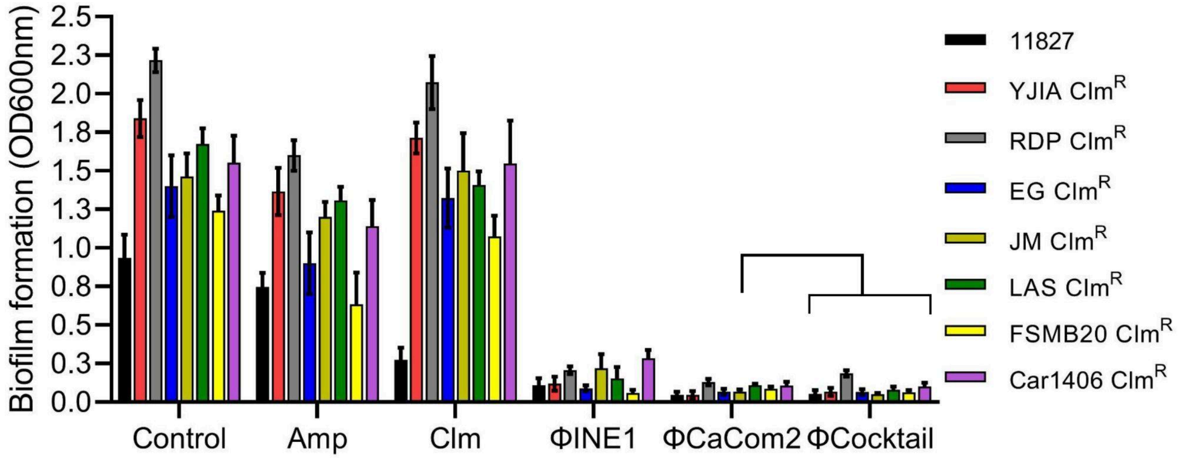


**Supplementary Figure S2.** Biofilm biomass quantified after 72 h of incubation using crystal violet staining. Treatments with ΦCaCom2 ($3 \times 10^8$ PFU/mL), ΦINE1 ($3 \times 10^8$ PFU/mL), and a phage cocktail (ΦCaCom2, ΦINE1, ΦCaSA2; each at $1 \times 10^8$ PFU/mL) were tested against seven clindamycin-resistant *C. acnes* isolates and compared to ampicillin (50 µg/mL) and clindamycin (30 µg/mL). Data are mean ± s.d. from three independent experiments ($n = 3$).

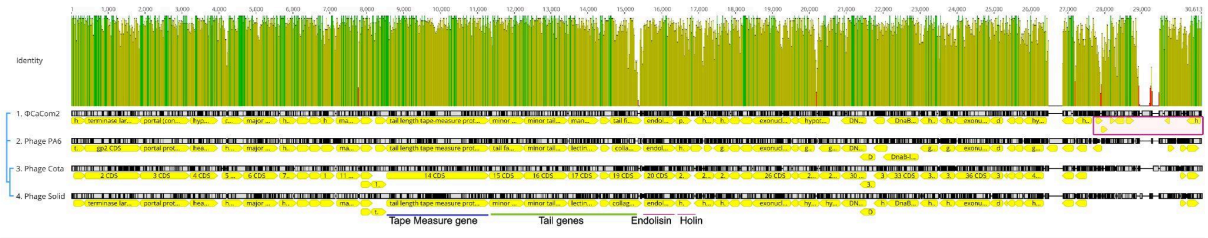


**Supplementary Figure S3.** Comparative genomic analysis of CaCom2 and previously characterized *C. acnes* phages (PA6, Cota, Solid, and Aquarius). Unique coding DNA sequences (CDSs) of phage CaCom2 are highlighted within a purple rectangle. The overall gene arrangement and GC content profiles were highly conserved among the five phage genomes. Whole-genome alignments were conducted using the Mauve plugin in Geneious Prime® 2025.1.

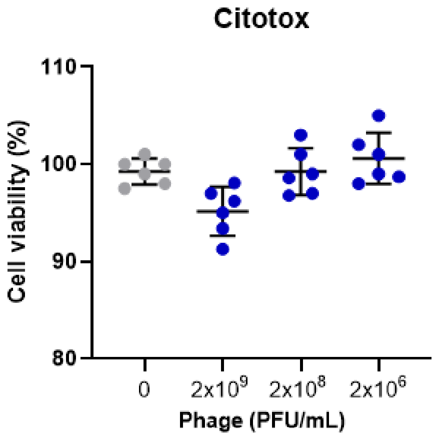


**Supplementary Figure S4.** The safety of the ΦCaCom2 phage formulation was investigated in human keratinocyte cultures. The effect of the phage on cell viability was evaluated in cultures exposed for 72 hours to high ($2x10^9$), medium ($2x10^8$), and low ($2x10^6$) concentrations of ΦCaCom2 phage. The data are presented as mean ± standard deviation (n=3 independent experiments).

### Supplementary tables.

**Supplementary table S1. Antibiotic susceptibility profile of *C. acnes isolates*.** The antibiotic susceptibility of the isolates was assessed using the disc diffusion assay, following the guidelines of the Clinical and Laboratory Standards Institute (CLSI) with minor modifications. The antibiotics tested included discs containing 10 µg ampicillin (AM), 30 µg tetracycline (TE), 30 µg minocycline (MH), 30 µg doxycycline (DO), 15 µg erythromycin (E) and 30 µg clindamycin (CLM). Antibiotic sensitivity is indicated by an "S" , and resistance is indicated by an "R".

| Strains | Species | AM | TE | MH | DO | E | CLM |
|---|---|---|---|---|---|---|---|
| ATCC 11827 | *C. acnes* | S | S | S | S | S | S |
| ATCC 6919 | *C. acnes* | S | S | S | S | S | S |
| DA-S | *C. acnes* | S | S | S | S | S | S |
| DA-B | *C. acnes* | S | S | S | S | S | S |
| VFP-Cut | *C. acnes* | S | S | S | S | S | S |
| Ca-JA19 | *C. acnes* | S | S | S | S | **R** | **R** |
| EG | *C. acnes* | S | S | S | S | S | **R** |
| USOG19A | *C. acnes* | S | S | S | S | S | S |
| YJIA | *C. acnes* | S | S | S | S | **R** | **R** |
| Jos-B1 | *C. acnes* | S | S | S | S | S | S |
| X1-Ca | *C. acnes* | S | S | S | S | S | S |
| MR | *C. acnes* | S | S | S | S | **R** | **R** |
| MP | *C. acnes* | S | S | S | S | S | S |
| JM | *C. acnes* | S | S | S | S | S | **R** |
| Chel | *C. acnes* | S | S | S | S | S | S |
| EB | *C. acnes* | S | S | S | S | S | S |
| RQ | *C. acnes* | S | S | S | S | S | S |
| Ca-YTJD | *C. acnes* | S | S | S | S | S | S |
| Ca-IA | *C. acnes* | S | S | S | S | S | S |
| RDP | *C. acnes* | S | S | S | S | **R** | **R** |

| | | | | | | | |
|---|---|---|---|---|---|---|---|
| Ca-VRL | *C. acnes* | S | S | S | S | S | S |
| LAS | *C. acnes* | S | S | S | S | S | **R** |
| FSMB20 | *C. acnes* | S | S | S | S | **R** | **R** |
| CAR1409 | *C. acnes* | S | S | S | S | **R** | **R** |

**Supplementary table S2. Antibiotic susceptibility profile of *C. acnes psedolysogens*.** The antibiotic susceptibility of pseudolysogens was assessed using the disc diffusion assay, following the guidelines of the Clinical and Laboratory Standards Institute (CLSI) with minor modifications. The antibiotics tested included discs containing 10 µg ampicillin (AM), 30 µg tetracycline (TE), 30 µg minocycline (MH), 30 µg doxycycline (DO), 15 µg erythromycin (E), 30 µg clindamycin (CLM). Antibiotic sensitivity is indicated by an "S" , and resistance is indicated by an "R".

| Parental Strains | Pseudolysogen | AM | TE | MH | DO | E | CLM |
|---|---|---|---|---|---|---|---|
| ATCC 11827 | PS1 | S | S | S | S | S | S |
| | PS2 | S | S | S | S | S | S |
| | PS3 | S | S | S | S | S | S |
| ATCC 6919 | PS6-3 | S | S | S | S | S | S |
| | PS6-7 | S | S | S | S | S | S |
| | PS6-11 | S | S | S | S | S | S |
| Ca-JA19 | PS-JA1-1 | S | S | S | S | S | S |
| | PS-JA1-2 | S | S | S | S | S | S |
| | PS-JA1-3 | S | S | S | S | S | S |
| EG | PS-EG1 | S | S | S | S | S | S |
| | PS-EG4 | S | S | S | S | S | S |
| YJIA | PS-Y1 | S | S | S | S | S | S |
| | PS-Y3 | S | S | S | S | S | S |
| | PS-Y5 | S | S | S | S | S | S |
| MR | PS-MR1 | S | S | S | S | S | S |
| | PS-MR3 | S | S | S | S | S | S |

| | | | | | | | |
|---|---|---|---|---|---|---|---|
| JM | PS-JM1 | S | S | S | S | S | S |
| | PS-JM2 | S | S | S | S | S | S |
| RDP | PS-RD1 | S | S | S | S | S | S |
| | PS-RD2 | S | S | S | S | S | S |
| CAR1409 | PS-CA1 | S | S | S | S | S | S |
| | PS-CA2 | S | S | S | S | S | S |
